# Vertical in-flight wind selectivity enables substantial energy savings in a migratory bird

**DOI:** 10.64898/2026.11.26.769459

**Authors:** Ance Priedniece, Viesturs Vīgants, Oskars Keišs, Ivo Dinsbergs, Māris Jaunzemis, Valts Jaunzemis, Elza Marija Zacmane, Raphaël Nussbaumer, Martins Briedis

## Abstract

Birds are known to change altitude during long migratory flights, yet drivers and benefits of this behaviour remain poorly understood. Here, we use 113 long-distance flights from 30 multi-sensor geolocator-tracked Common Starlings *Sturnus vulgaris* to test whether altitude changes are associated with vertical wind heterogeneity and reflect selective use of winds to reduce energy costs. Birds adjusted altitude in response to vertical wind structure, exhibiting high wind selectivity under headwinds and strong vertical wind heterogeneity, but showed low selectivity under tailwinds or uniform wind conditions. High wind selectivity substantially reduced flight costs, allowing birds to save >20% of flight costs during autumn migration compared to simulated altitude use, whereas low selectivity incurred no additional costs during spring migration dominated by tailwinds. Our findings reveal that a small passerine migrant actively navigates the aeroscape to reduce metabolic flight costs, highlighting a behavioural mechanism with broad implications for migratory performance and survival.

## Introduction

Each year, billions of birds fly thousands of kilometres to carry out their seasonal migrations from breeding to non-breeding grounds and back again. While flight is the most efficient form of animal locomotion in terms of distance travelled per unit of time (Newton 2023), it is also among the most energy-intensive forms of movement, with flapping flight requiring eight to nine times more energy than resting (Winkler et al. 2016). Thus, carrying out these large-scale movements requires substantial amount of energy. Despite this, migratory species are capable of exceptionally long nonstop flights that can last multiple days (e.g., Dufour et al. 2025; Gill et al. 2009). To meet the energetic demands of these highly costly journeys, birds can combine two complementary strategies: (1) they may build up fat reserves prior to initiating migratory flights and (2) reduce energy expenditure during the flight itself by travelling under favourable atmospheric conditions, particularly winds.

Accumulation of fat reserves before migration has been widely documented across avian taxa (Berthold 2001; Guglielmo 2018; McWilliams et al. 2004). As fat is the primary fuel for migratory flight (Guglielmo 2018), its energy content and implications for flight performance have been extensively studied (Hedenström & Alerstam 1997; Newton 2023; Pennycuick 2008; Winkler et al. 2016). Although atmospheric conditions have been shown to substantially influence bird and bat flight costs (Scacco et al. 2026), the extent to which birds can reduce the energetic costs of flight through behavioural adjustments while airborne remains unclear.

Studies on departure decisions show that birds tend to initiate migratory flights under more favourable weather conditions (Åkesson & Hedenström 2000; Battley 1997; Gill et al. 2009; Pyle et al. 1993; Richardson 1990a; Schaub et al. 2004), while broad-scale radar studies show that birds preferentially fly at altitudes with more favourable wind conditions (Dokter et al. 2013; Horton et al. 2016; Kemp et al. 2013; Liechti & Schmaljohann 2007; Mateos & Liechti 2012). While these studies have substantially advanced our understanding of how weather influences bird migration, they do not capture altitude dynamics throughout entire flights. More recent high-resolution tracking studies show that birds frequently adjust flight altitude during migratory flights, often by hundreds or even thousands of meters (Dufour et al. 2025; Sjöberg et al. 2021). These changes can occur irregularly (Bowlin et al. 2015) or coincide with transitions between day and night, as observed in Great Reed Warblers *Acrocephalus arundinaceus* (Sjöberg et al. 2021). While the causes and potential benefits of these altitudinal shifts remain unclear, one hypothesis is that they are driven by variation in atmospheric conditions, including ambient temperature, solar radiation, wind, relative humidity and air density (Senner et al. 2018; Sjöberg et al. 2021).

In seabirds, shearwaters exhibit varying degrees of wind selectivity during foraging flights by adjusting their heading according to prevailing wind conditions, thereby enabling energy- and time-efficient flight (Harris et al. 2025). They showed low wind selectivity during target-oriented flights and under low wind speeds but became highly selective under strong crosswinds and tailwinds that influenced flight direction.

These findings are intriguing in the context of the frequent altitude changes observed during migratory flights in a several species (Dufour et al. 2025; Lindström et al. 2021; Sjöberg et al. 2021), as similar principles may guide in-flight decisions along the altitudinal axis.

Liechti (2006) suggests that wind conditions experienced aloft can significantly affect flight costs, potentially allowing birds to double or halve their cost of transport. Importantly, wind conditions are highly heterogeneous both horizontally and vertically along migratory routes, with wind direction and speed varying substantially (Dokter et al. 2013; Liechti 2006). Consequently, birds would need to search and adaptively select flight altitudes with profitable wind conditions throughout the flight to fully capitalize on these potential benefits. Although climbing and sinking during a flight inevitably incurs additional energetic costs (Nudds & Bryant 2000), they are relatively negligible compared to the costs of forward flight (Brodersen et al. 2026; Norevik et al. 2021). Despite the substantial theoretical advantages of prospecting and selecting favourable wind conditions, empirical quantification of wind selectivity across altitudes over the course of migratory flights and its associated energetic consequences remains limited.

In this study, we examine migratory flight altitudes of a short-distance migrant, the Common Starling *Sturnus vulgaris*. Starlings rely on the most energy-demanding form of flight – flapping flight – to power their migration (Tobalske 1995). High wind selectivity may therefore be crucial for minimizing energy expenditure during migration. Consequently, we hypothesize that flight altitude changes observed in geolocator-tracked Starlings are related to wind conditions and result in high wind selectivity relative to overall windscape. In particular, we expect wind selectivity to be high during flights with strong headwinds and large differences in wind conditions across altitudes, where altitude selection can yield substantial energetic benefits. Conversely, wind selectivity should be low under tailwinds and uniform wind conditions across altitudes. Our study demonstrates how wind-related altitude adjustments during flight enable substantial energy savings in a short-distance migrant and a workable framework to evaluate the importance of this behaviour across species and migratory strategies.

## Materials and methods

### Data collection

In 2020, we established two field sites: one with 200 nest boxes (100 + 100 at two nearby locations) in western Latvia around Lake Engure (57.27°N, 23.11°E) and one with 100 nest boxes in eastern Latvia in Zeltaleja (57.19°N, 26.86°E). The two sites are approx. 230 km apart. Between 2020 and 2023, we captured breeding adult birds, mainly with traps inside nest boxes, and equipped them with multi-sensor data loggers, i.e., geolocators (SOI-GDL3pam, Swiss Ornithological Institute; Liechti et al. 2018), following the field protocol described in Vīgants et al. (2023). These geolocators weighed 1.96 ± 0.08 g (SD) including the harness, which represented 2.5% of the Starling’s average body weight (79.2 ± 3.9 g, n = 500) and thus, comply with recommended weight threshold for biologging device attachment (Barron et al. 2010; Brlík et al. 2020). In total, we tagged 308 Starlings with geolocators attached to the back of the birds using a non-flexible nylon leg-loop harness. The devices were programmed to measure ambient temperature every 30 minutes, light intensity and relative acceleration every 5 minutes, and atmospheric pressure every 30 minutes in loggers deployed in 2020-2021 and every 5 minutes in those deployed in 2022-2023. In total, 47 previously deployed geolocators were recovered, including six recovered two years after deployment, resulting in an overall recovery rate of 15.6% (excluding eight loggers lost between consecutive captures). Consistent with an earlier assessment of logger effects in our study population (Vīgants et al. 2023), we did not find significant negative effects of geolocators on adult return rates compared with ringed-only birds (tagged: 47/301 = 15.6%, ringed-only: 56/288 = 19.4%; χ² = 1.496, p > 0.05).

Data from 31 of the 46 retrieved geolocators contained information from at least one migratory season and were thus usable for analysis. The remaining geolocators could not be used due to sensor malfunction or premature battery failure caused by physical damage. Four of the 31 usable geolocators recorded data from a single migratory season (autumn), whereas three were recovered two years after deployment and contained data of more than one migration cycle.

### Data analyses

The data were analysed in R v.4.4.1 (R Core Team 2024) using the package *GeoPressureR* v.3.2.0 (Nussbaumer & Gravey 2024) to reconstruct migratory tracks from raw geolocator data, following the workflow outlined in Nussbaumer & Nussbaumer (2024). In short, *GeoPressureR* combines an observation model based on pressure and light data together with a movement model within hidden Markov model to estimate the most likely migration trajectory. The pressure observation model matches the pressure timeseries against the ERA5 atmospheric reanalysis (Muñoz Sabater 2019), while the light data is converted to solar zenith angles to produce a likelihood at each candidate location. The movement model uses accelerometer-derived flight periods and their durations together with ERA5 wind data to estimate the airspeed required to travel between successive candidate stopover sites. Importantly, the model favours energetically plausible airspeeds when evaluating candidate paths (Nussbaumer et al. 2023a), which typically fall near the theoretical maximum range speed – the airspeed that maximizes distance travelled per unit of energy (Pennycuick 1969) and is expected under an energy-minimizing migration strategy (Alerstam & Lindström 1990). Hidden Markov model then combines both observation models to identify the sequence of stopover sites connected by realistic airspeeds, yielding a migration trajectory with typical uncertainty of <100 km (Nussbaumer et al. 2023a). For further analyses, we used this most likely path, the single trajectory that best fits the whole sequence of observations and flights, as the bird’s most probable migration route.

Based on the reconstructed migration trajectory, we focused exclusively on migratory flights with a minimum duration of six hours, as shorter flights offer limited opportunity for altitude exploration. This threshold ensured a minimum of 11 altitude measurements per flight (excluding take-off and landing) at 30-min recording interval, allowing us to capture potential altitude adjustments. This resulted in a total of 113 migratory flights from 30 individuals, four of which were tracked over two annual cycles. Of these, 64 flights (57%) were recorded during autumn migration (June 1^st^ – December 31^st^), while the remaining 49 flights (43%) occurred during spring migration (January 1^st^ – May 31^st^). The duration of the analysed flights ranged from 6 h 10 min to 24 h 30 min (mean ± SD = 11 h 5 min ± 3 h 50 min).

Flight altitude and day-night classification were derived using the *pressurepath_create* workflow in *GeoPressureR*: altitude was computed from geolocator-recorded pressure by referencing ERA5 surface pressure, temperature, and surface elevation, and day-night status was determined from local sunrise and sunset times calculated with *suncalc* algorithm (Nussbaumer & Nussbaumer 2024). To account for seasonal and latitudinal variation in day/night length, we normalized the timestamp of each pressure measurement relative to local sunrise and sunset. We refer to this normalized value as the normalized time index (NTI).

Further, we used a generalised additive mixed-effects model (GAMM; function *gamm4* in the *gamm4* package; Wood et al. 2026) to quantify flight altitude changes throughout the diel cycle and compare altitude patterns between migratory seasons. The response variable (altitude) was Yeo-Johnson transformed (λ = 0.64, estimated from the data) to improve normality and homoscedasticity. Season was included in the model as a fixed factor (*fseason*), and we allowed a season-specific smooth of NTI using a factor–smooth interaction. Random intercepts were included for individuals (*@ird_ID*) and for flights nested within individuals (*flight_ID* within *@ird_ID*) to account for inter-individual variability and repeated measurements within flights.

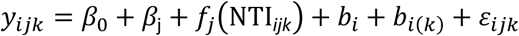

where *y_ijk_* is the Yeo-Johnson transformed altitude, *βj* is the fixed effect of season *j*, *fj* (NTI*_ijk_*) is the season-specific smooth function of normalized time index, *bi∼N(0,σ^2^bird)* is the random intercept for bird *i*, *bi(k)∼N(0,σ^2^flight)* is the random intercept for flight *k* nested within bird *i*, and *εijk∼N(0,σ^2^)* represents the residual error.

Model predictions were back-transformed to metres. Seasonal differences in the diel pattern were tested using an equivalent model in the *mgcv* package (Wood 2017) with first-order autoregressive residual errors.

Wind selectivity

To quantify how effectively Starlings exploit optimal wind conditions across altitudinal levels during migratory flights, we developed a Wind Selectivity Index (WSI) for each migratory flight. This index quantifies the extent to which birds choose energetically favourable flight altitudes relative to the range of wind conditions available at any given time during the flight.

For each flight, timestep (either 30-min for 2020-2021 tracked individuals or 5-min for 2022-2023 tracked individuals), and location along the horizontal flight path, we calculated the wind support experienced by the birds using the *windsupport* function in the R-package *GeoPressureR* (Nussbaumer et al. 2023a, b). Additionally, we obtained wind data (u and v components) from the pressure-level ERA5 dataset (Hersbach et al. 2023) for all 11 pressure levels between 1000–750 hPa at 25 hPa intervals, corresponding roughly to altitudinal levels from ground level up to 2500 m above sea level and encompassing the range of flight altitudes recorded in Starlings.

Selectivity for most favourable wind conditions during a migratory flight may not solely involve finding the flight altitude with maximal tailwinds (or minimal headwinds) at each timestep. This is because changes in flight altitude, particularly climbs, incur additional energetic costs (Nudds & Bryant 2000). To account for this, wind selectivity analyses were based on total energy expenditure associated with both vertical and horizontal movement. Accordingly, the optimal altitudinal path during a migratory flight was defined as the path minimizing overall energy requirements given wind support and altitude changes. We used the R-package *afpt* (Klein Heerenbrink 2023) and its function *findMaximumRangeSpeed* to estimate flight costs for each 30-min timestep across all considered altitudinal levels throughout a migratory flight. Estimates incorporated bird morphological parameters (body mass = 79.2 g, the mean in our dataset; wingspan = 38 cm; wing area = 230 cm^2^; Bruderer et al. 2010), altitude-specific wind speed and direction, and climb rate required to reach each altitudinal level from the previous level. Consequently, the WSI for each flight was calculated as follows:

1. For each timestamp and location along the migration path, we estimated wind support across all available altitude levels relative to the bird position, flight direction of the bird as revealed from geolocator tracks.
2. For each flight segment (30-min interval, data from geolocators with 5-min sampling rate were downscaled) and altitudinal level, we calculated all possible climb angles in degrees between the focal altitude level and all others based on the distance travelled during that timestep (groundspeed of the bird x time) and altitude difference between the layers.
3. Using *findMaximumRangeSpeed* function from the R-package *afpt* (Klein Heerenbrink 2023), we produced flight cost matrix (kJ/km) for all possible altitude change scenarios based on wind speed, wind direction, and climb rate.
4. We estimated energetic costs (kJ/km) of each migratory flight of the tracked birds by drawing the recorded flight altitude path through the flight cost matrix.
5. Null model: Because optimal wind conditions are often unevenly distributed across both horizontal and vertical dimensions of the flight environment, we generated 10 000 alternative flight altitude paths for each flight based on realistic altitude change characteristics of Starling flights observed in our data. Specifically, the simulated paths always started and ended at the same altitude as the recorded flights, while altitude changes between consecutive timesteps were generated using the following parameters: altitude change probability = 0.5, magnitude of altitude change = 1 ± 2 altitude levels per change event (mean ± SD).
6. For each of the generated flight altitude paths, we calculated energetic costs (kJ/km) and compared them to the estimated energetic costs of the recorded bird path.
7. Finally, the WSI for the bird’s flight path was calculated as quantile of the energetic costs of bird flight path relative to the energetic costs of the 10 000 alternative paths.

The resulting WSIs of bird’s path range from 0 to 1, where WSI = 0 indicates the worst possible choice with the highest flight costs and WSI = 1 indicates perfect selectivity with minimal flight costs per km travelled. Correspondingly, WSI values close to 0.5 would be attained following the null model and thus can indicate lack of selectivity.

To test if the observed WSIs of Starling flights were significantly higher than 0.5 – indicating that birds selected flight altitudes with lower energetic costs than expected under random altitude choice – and whether this differed between migration seasons, we fitted a linear mixed-effects model with migration season as explanatory variable. We included bird ID as a random factor to account for possible individual variation (e.g., due to experience, body condition) in wind selectivity. To test whether wind selectivity was more pronounced under certain wind conditions, e.g., under headwinds or tailwinds, we employed a linear mixed-effects model with the average available wind support (per flight) and season as fixed effects and bird ID as a random effect. All models were fitted using the *lmer* function from the *lme4* R-package (Bates et al., 2015). Pairwise comparisons between factors were performed using estimated marginal means (EMMs) with the *emmeans* R-package (Lenth, 2024).

Further, we modelled energy savings as a function of wind support using a nonlinear mixed-effects exponential decay function:

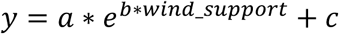

where *a* (amplitude), *b* (decay rate), and *c* (asymptote) were fixed effects, with *a* incorporating random effects across individuals to account for variability in energy savings. The model was fitted via maximum likelihood estimation using the *nlme* R-package (Pinheiro & Bates, 2000), capturing continuous decay dynamics from headwind to tailwind conditions while accommodating individual differences in response magnitude.

Lastly, we modelled the relationship between wind selectivity index (WSI) and energy savings using a nonlinear mixed-effects sigmoidal function:

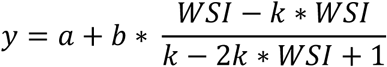

where *a* (baseline), *b* (amplitude), and *k* (curvature) were fixed effects, with *a* including random effects across individuals to account for inter-individual variability. The model was fitted via maximum likelihood estimation using R-package *nlme*, capturing both population-level dynamics and individual differences in baseline energy savings.

Finally, we expressed the energy savings in g of body fat assuming a standard energy content of 39.54 kJ g^-1^ of body fat in birds (Berthold 2001; Carey 1996).

## Results

Most of the tracked Starlings wintered in the British Isles, while two individuals travelled shorter distances and wintered in Denmark and the Netherlands, whereas one bird wintered on the east coast of Spain (**Figure 1a**). Autumn migration took place between end of September and early December, while spring migration from late February to early April **(Figure 1b).** A few individuals engaged in moult-migration that took place between mid-June and mid-July.

**Figure 1.**
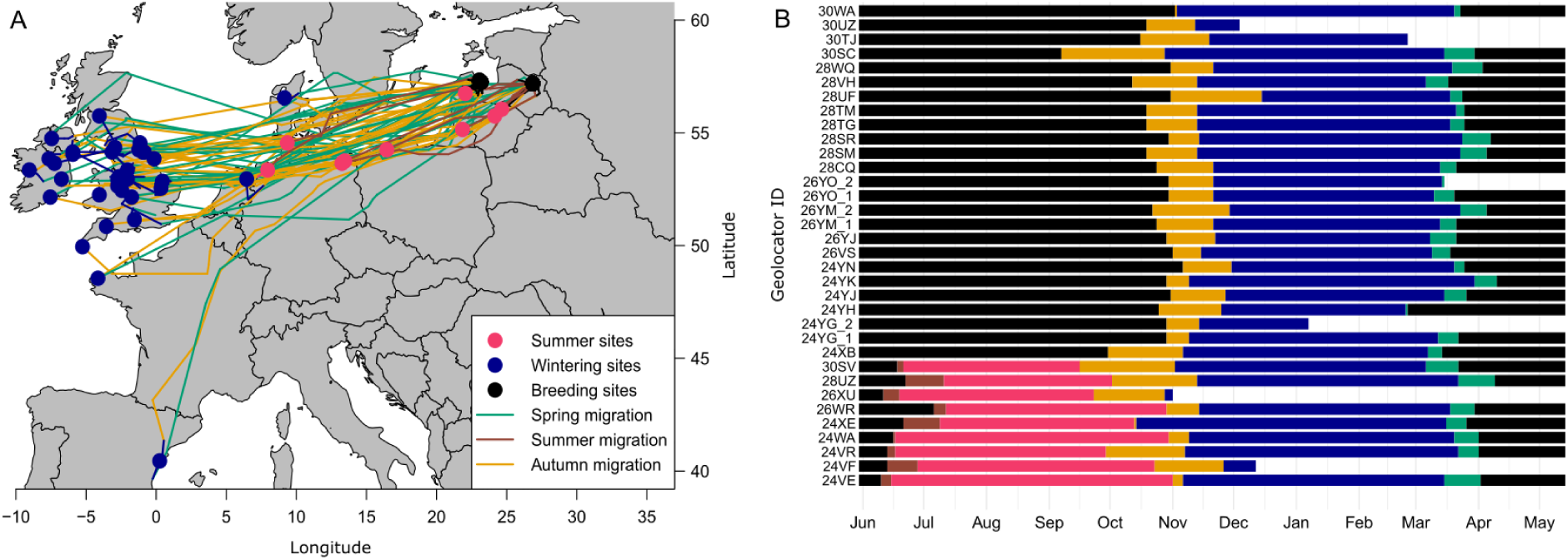
(A) Migratory tracks and (B) the timeline of the annual cycle of Starlings breeding in Latvia. The colours in the map legend also refer to the timeline. Shortened timelines are from geolocators that stopped recording early. Geolocator IDs 24YG_1/24YG_2, 26YM_1/26YM_2, 26YO_1/26YO_2, and 28UZ/30SC represent two migration cycles of the same individual.

### Flight patterns

Starling migratory flights were predominantly nocturnal. Most flights started around sunset and finished around sunrise, with some extending into the day (**Figure 2**). Average flight altitudes followed a non-linear pattern throughout the night, with the highest altitudes recorded shortly after sunset, followed by a gradual decrease until landing. In autumn, altitudes formed an additional, smaller peak after midnight that was not observed in spring. Diurnal flights were mostly carried out at lower altitudes than nocturnal flights in both seasons. Migratory flight altitudes were significantly higher in spring than in autumn (paired Wilcoxon signed-rank test: V_24_ = 270, p < 0.001) and associated with steeper ascent at sunset and steeper descent before midnight in spring (edf = 5.9, F = 11.6, p < 0.001).

**Figure 2.**
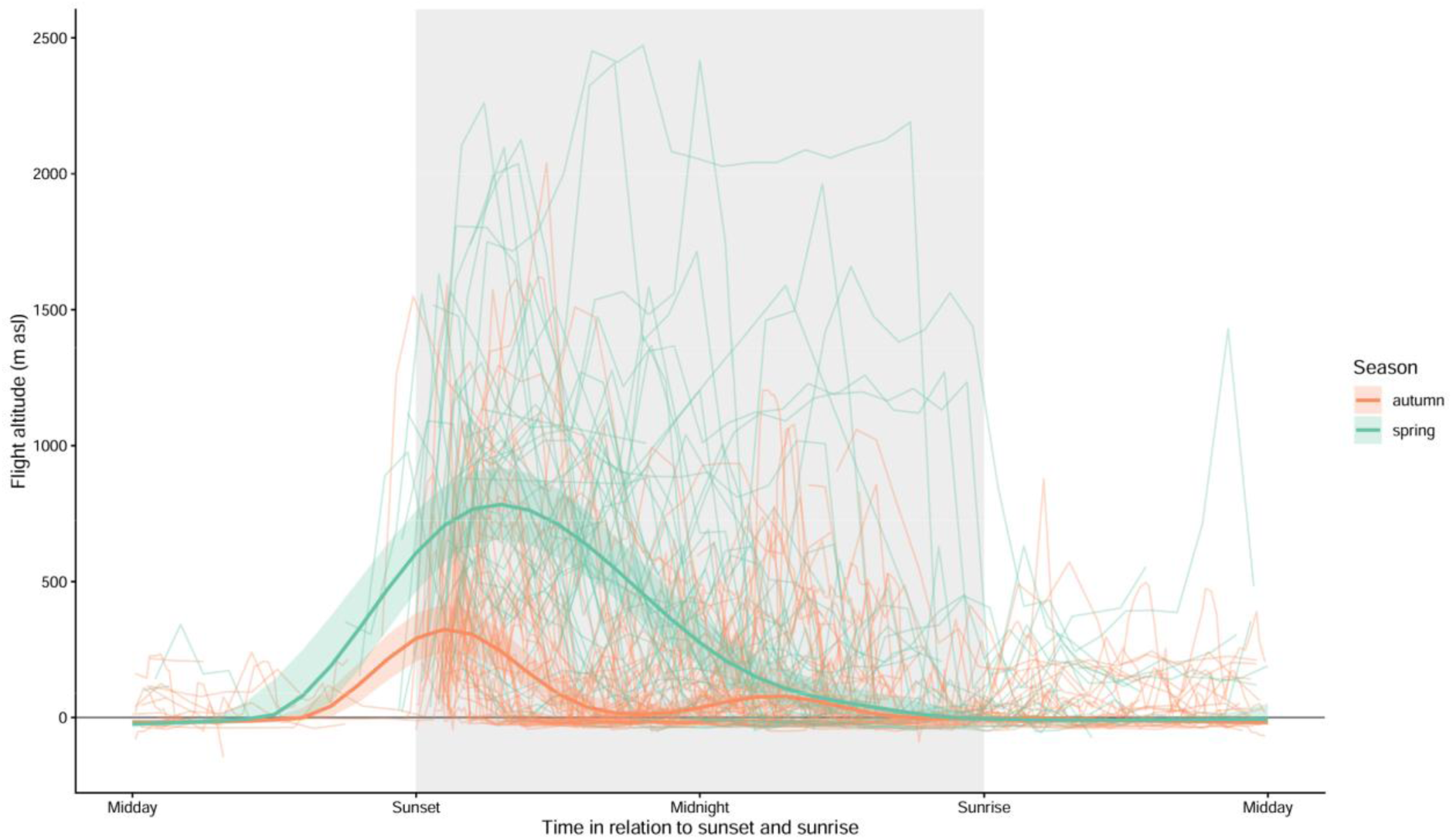
Migratory flight altitudes in relation to sunset and sunrise. Flights with a duration of at least six hours were used. Thin lines represent individual flights, bold line and ribbon represent the population level prediction of the GAMM model and its 95% confidence interval. Flight altitudes below zero are an artefact caused by specific atmospheric conditions and correspond to flight altitudes near sea level.

### Wind selectivity

Available wind conditions in autumn and spring were significantly different from one another along the migratory tracks of Starlings with prevailing headwinds in autumn (mean wind support ± SD = −8.5 ± 19 km/h) and strong tailwinds in spring (25.4 ± 20.8 km/h; β = −33.9 ± 3.6 SE, t = −9.5, p < 0.001). Such wind conditions correspond to flight energy expenditure of 0.795 kJ/km in autumn and 0.366 kJ/km in spring (not considering any altitudinal climbs during the flight) when Starlings are flying at maximum range airspeed – the speed at which a bird can travel the farthest distance for a given amount of energy (autumn = 41.4 km/h, spring = 33.4 km/h). Consequently, birds would require a total of 1591 kJ of energy to complete the approx. 2000 km long autumn migration and a total of 730 kJ of energy for the same length return journey in spring. This corresponds to a total of 40.2 g and 18.5 g of body fat required to fuel the two migrations, respectively, assuming that fat is the sole fuel source and has an energetic content of 39.54 kJ g^-1^ in birds (Berthold 2001; Carey 1996).

The mean wind selectivity index (WSI) of Starling flights during autumn migration was 0.82 (95% CI: 0.74–0.89), significantly higher than the null expectation of 0.5 (t_62_ = 8.35, p < 0.0001), indicating strong selectivity for optimal wind conditions. In spring, however, the mean WSI of Starling flights was 0.5 (95% CI: 0.41–0.58) which did not differ from the null model (t_63_ = −0.1, p = 0.92; **Figure 3a**), indicating lack of selectivity. Overall, the WSI was negatively related to the mean available wind support during the flights, so that high WSIs were typical for flights dominated by headwinds while low WSIs were observed during flights dominated by tailwinds (β = −0.005 ± 0.001 SE, t = −3.6, p < 0.001; **Figure 3b**). Thus, in-flight altitude changes during migratory flights were associated with prevailing wind conditions, particularly reflecting avoidance of headwinds (see **Figure 4** for an example of a tracked Starling flight and the corresponding wind conditions during it).

**Figure 3.**
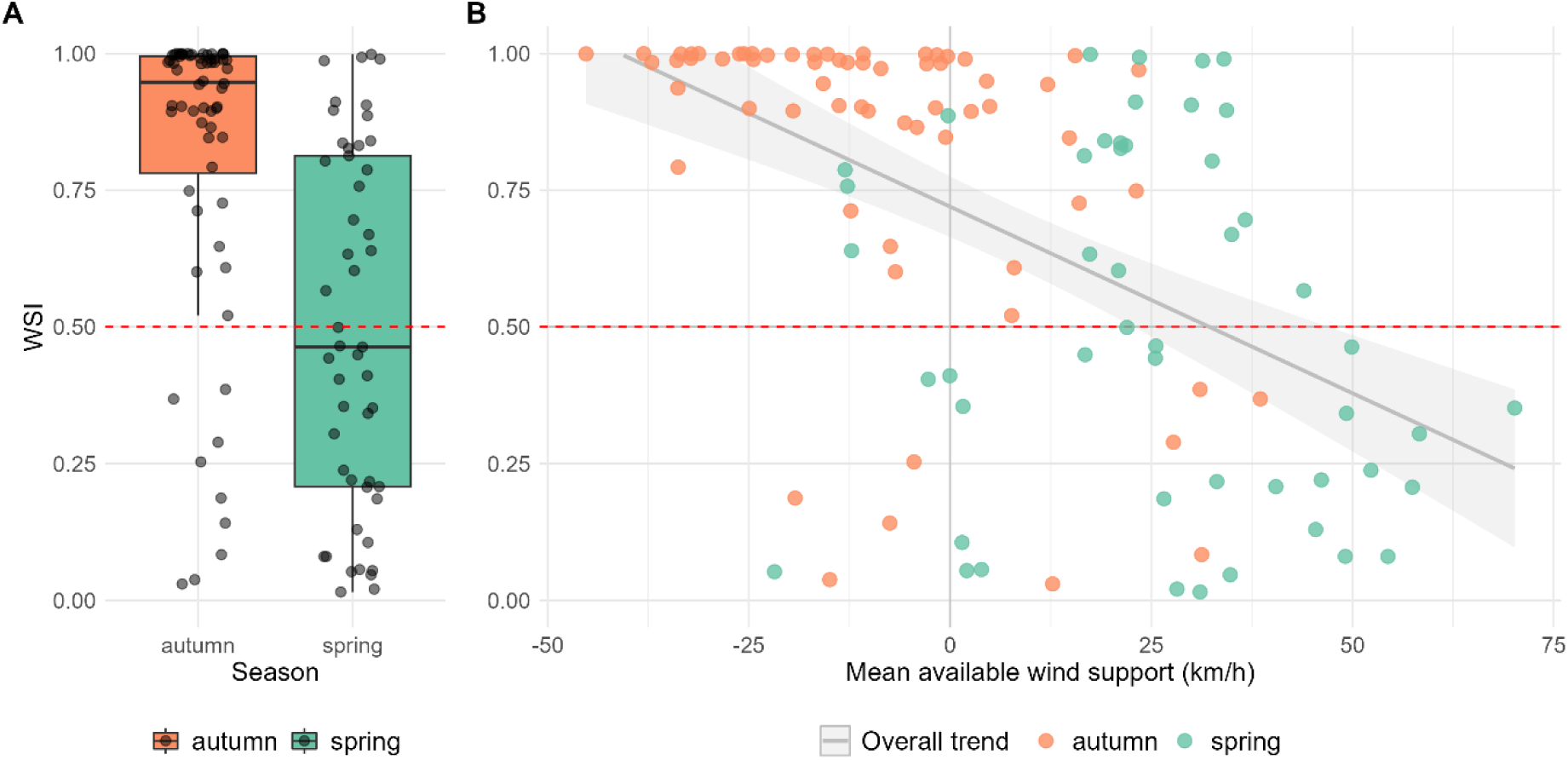
Wind selectivity of Starlings during long (> 6 h) migratory flights. (A) Wind selectivity index (WSI) of individual flights depending on migration season. Horizontal lines inside boxes depict median values, boxes show IQRs, whiskers extend 1.5 times the IQR, and points denote individual flight data. (B) Relationship between WSI and the mean available wind support during each flight (negative values correspond to headwind, positive values to tailwind). The line shows predicted model fit along with 95% CI.

**Figure 4.**
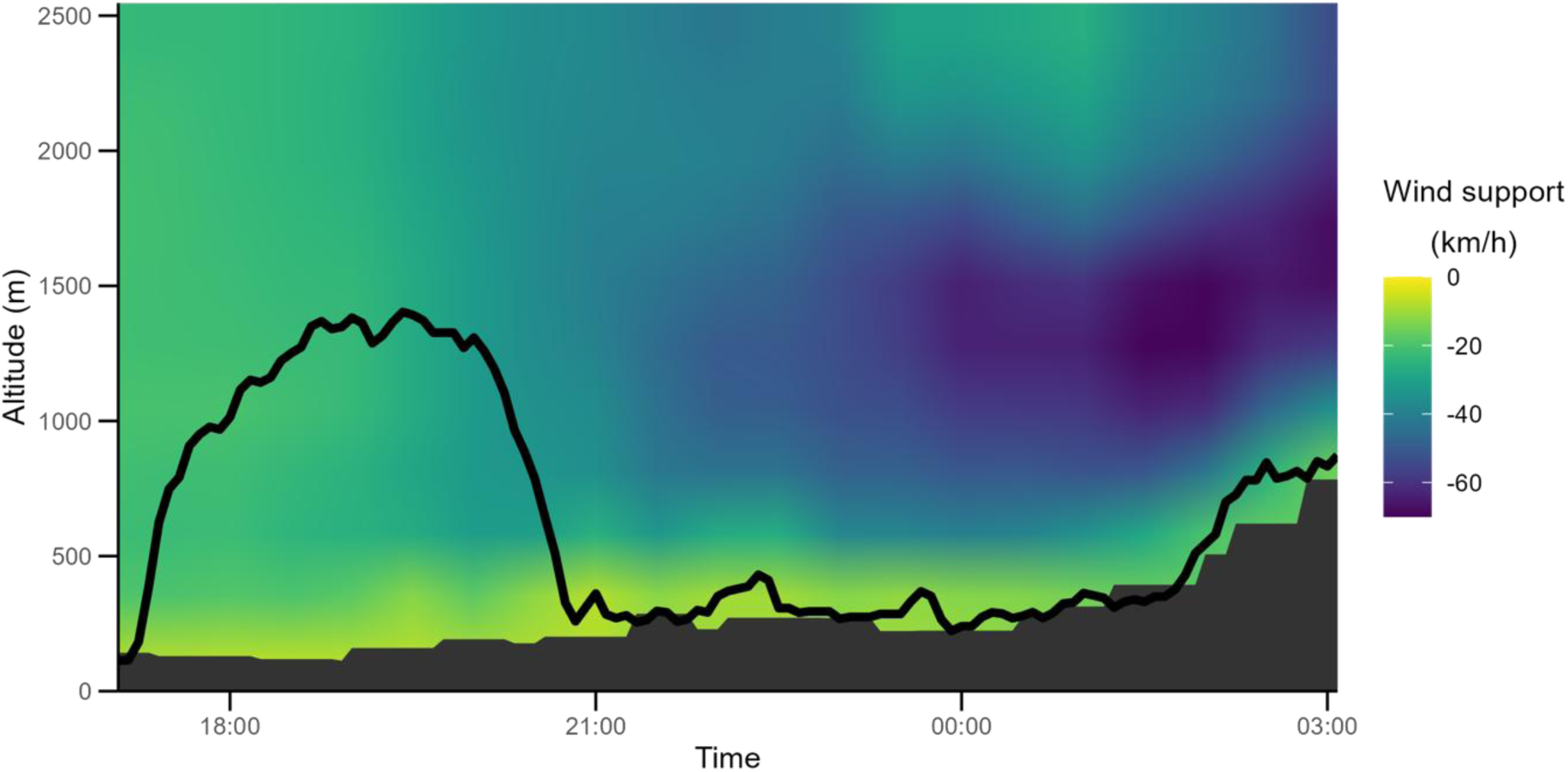
An example of wind conditions during a migratory flight of a Starling 30SV. Wind selectivity index (WSI) for this flight was 0.94. The flight lasted for 10 hours between 24–25 October. The dark polygon on the bottom of the figure denotes the average ground elevation and the black line depicts the flight path. Background colours indicate the strength of wind support at different altitudes during the flight, where negative values correspond to headwind. Notice the altitude drop around 20:30h that coincides with increased headwinds ahead (blue background). See Supporting Information for figures of all 113 analysed migratory flights.

Energy savings showed a strong sigmoidal relationship with WSI. High WSI values were strongly associated with increased energy savings while energy savings at low WSI values were not significantly different from 0 (**Figure 5b**; a = 0.02, p = 0.406). Rapid increase in energy savings occurred at WSI > approx. 0.9 (k = 0.96, p < 0.001). At WSI = 1, the model predicted energy savings of 0.497 kJ km^-1^. This pattern is expected because both WSI and energy savings depend on how much flight costs vary among altitudes: when costs are similar, even choosing the best altitude saves little energy, whereas larger differences in costs offer greater potential savings. Consistent with this, low WSIs were mostly observed during flights with relatively uniform wind conditions across altitudes, producing a narrow window of flight costs and minimal energetic penalties for sub-optimal altitude choice. High WSIs in contrast often occurred during flights with large vertical variation in wind conditions, creating a broad window of possible flight costs and considerable scope for energy savings. As a result, Starlings achieved meaningful energy savings primarily during flights dominated by headwinds, whereas flights with neutral or tailwind support offered little opportunity for savings (**Figure 5a**). Energy savings followed an exponential decay relationship with mean available wind support during flights. The nonlinear mixed-effects model showed significant amplitude (a = 0.022 ± 0.007 kJ km^-1^, p = 0.002) and decay rate (b = −0.084 ± 0.007 kJ km^-1^, p < 0.001), confirming maximal energy savings under headwind conditions. Under tailwinds, predicted savings approached an asymptote that did not differ from zero (C = −0.002 ± 0.017 kJ km^-1^, p = 0.925).

**Figure 5.**
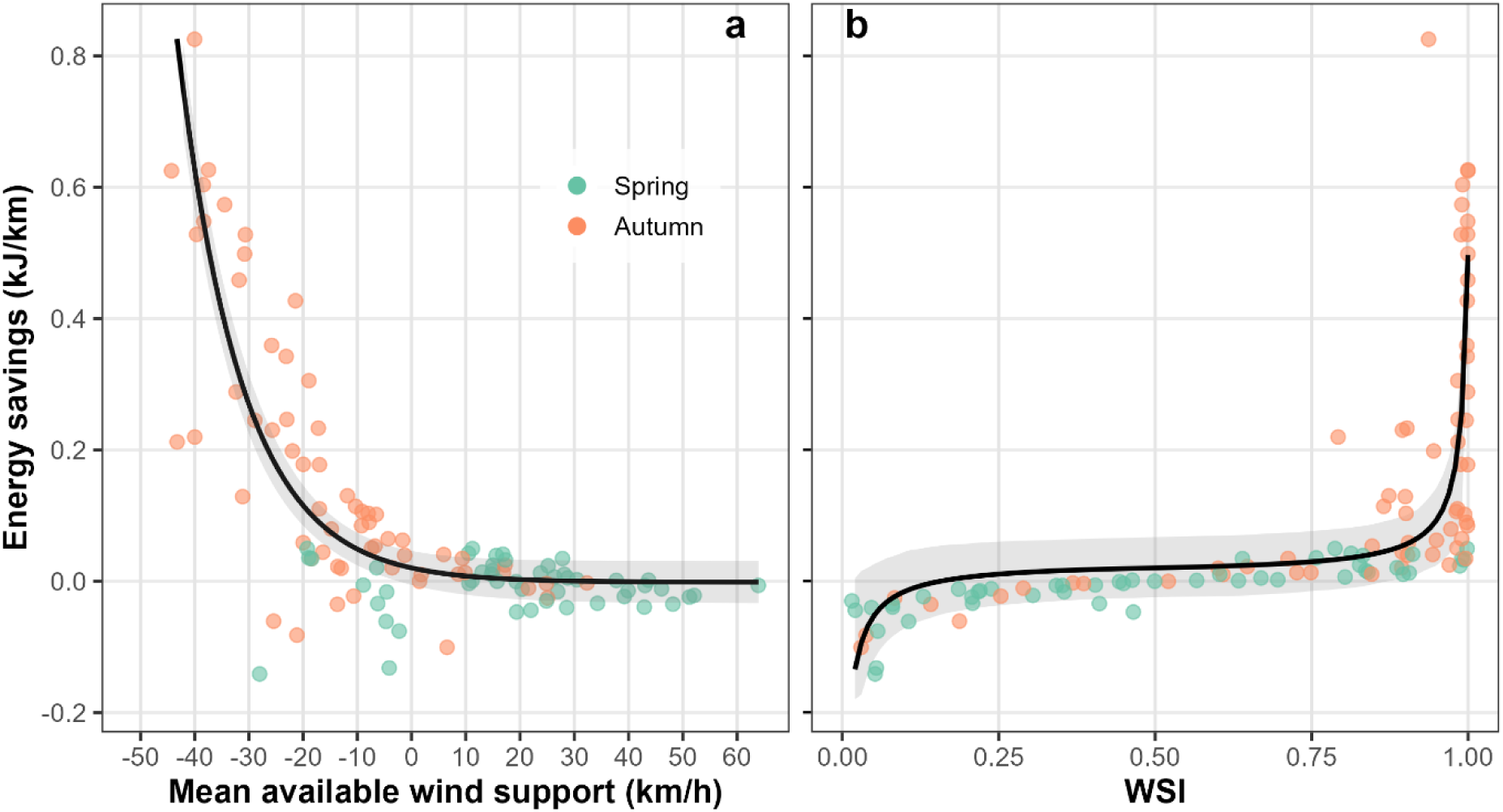
Energy savings (kJ/km; in comparison to modelled flight altitude paths with WSI = 0.5) achieved by Starlings during migratory flights as a function of (a) mean available wind support and (b) wind selectivity index (WSI). Each point represents an individual flight during spring (green) or autumn (orange) migration. Lines show predicted model fits from nonlinear mixed-effects models, with the shaded areas indicating the 95% CIs. Negative wind support values in (a) correspond to headwinds, while positive values indicate tailwinds. Note the increase in energy savings with increasing headwinds and high WSIs, and near-zero energy savings/losses under tailwinds and low WSIs.

The observed wind selectivity during autumn migration would enable birds to conserve 431 ± 639 kJ (SD) of cumulative flight energy expenditure, equivalent to 10.9 ± 16.2 g of body fat. In contrast, the absence of wind selectivity during spring migration would require birds to expend only additional 15 ± 79 kJ or 0.4 ± 2 g of body fat on average to complete the approx. 2000 km long one-way migratory journey. This corresponds to 22.4% energy savings in autumn, and 1.6% energy expenditure increase in spring in comparison to the estimated flight costs for modelled flight altitudes.

## Discussion

We found that migratory Starlings frequently adjust their flight altitude during migratory flights, and these altitude shifts are often related to wind conditions. Wind-related altitude shifts were most pronounced under headwinds when birds showed high wind selectivity (higher WSI). During autumn migration, high wind selectivity reduced the total flight costs required to complete the journey by more than 20% compared to non-selective altitude use.

In contrast, during spring migration, when birds encountered tailwinds more often, lack of wind selectivity did not incur additional energetic costs. Seasonal differences in wind selectivity were likely driven by contrasting wind conditions, with prevailing westerly winds along the migration routes (Hurrell & Deser 2010) acting as headwinds in autumn and tailwinds in spring. Low wind selectivity was often observed during flights that had relatively similar wind conditions across altitudes and hence a narrow window of flight costs in which choosing suboptimal flight altitudes incurred minor energetic penalties. High wind selectivity, on the other hand, was frequently observed during flights that had a broad window of flight costs and hence considerable scope for energy savings through in-flight altitude adjustments.

Tracked Starlings did not consistently choose the energetically most optimal flight altitudes that would allow covering largest distance at the lowest costs. They showed clear avoidance not only of strong headwinds but also strong tailwinds, consistent with observations of Able (1970), indicating that extreme wind conditions – regardless of direction – were unsuitable for sustained migratory flight. In some flights, headwinds at certain altitudes approached the species’ maximum airspeed (∼61.2 km/h; Bruderer and Boldt 2001), rendering forward movement impossible. Likewise, very strong tailwinds may be avoided because they can impair flight control or stability, given that higher wind speeds are often accompanied by increased turbulence (Stull 2015). Strong winds may also put birds at risk of being blown off course and impair their ability to compensate for wind drift or locate suitable landing/stopover habitats (Kemp et al. 2013).

The failure to consistently select the energetically most optimal altitude may reflect the influence of other factors, such as temperature, humidity, solar radiation, predation, and orientation (Dufour et al. 2025; Lindström et al. 2021; Senner et al. 2018; Sjöberg et al. 2021, 2023), as well as how birds assess available wind conditions during flight. Starlings tended to explore winds in upper altitude layers early in the flight and descend soon after conditions aloft deteriorated. However, they generally did not repeat these exploratory climbs later in the flight even when near-ground conditions became unfavourable. Such early-flight sampling of wind conditions at different altitudes has also been documented in nocturnal migrants during the first two hours after sunset (Bruderer et al. 1995). Migrant birds have likewise been shown to reach peak altitudes during the first 30% of the night before gradually descending thereafter (Horton et al. 2016). The smaller secondary peak after midnight observed only in autumn could reflect additional exploration later in some flights, potentially in response to unfavourable winds. Exploratory ascents throughout the flight have also been reported in nightjars, although they have not been explicitly linked to wind conditions (Norevik et al. 2021).

The varying strength of wind selectivity we observed may reflect the pattern described by Kemp et al. (2013), who reported that birds generally flew at the lowest altitude that still provided tolerable wind conditions. Gauthreaux (1991) similarly found that birds flying at *higher-than-usual* altitudes positioned themselves at the lowest altitude where winds remained tolerable. This behaviour could reflect inability of birds to assess wind conditions at higher altitudes before ascending into them (Dokter et al. 2013; Liechti 2006), and that climbing itself incurs additional energetic costs (Liechti 2006; Pennycuick 2008), although these costs appear to be relatively small in the broader migratory flight cost context (Brodersen et al. 2026; Norevik et al. 2021).

Birds’ unwillingness to climb to altitudes with optimal wind conditions may also reflect other environmental constraints. At higher altitudes, reduced air density decreases lift and lower oxygen partial pressure limits aerobic capacity, requiring more metabolic power to stay aloft (Pennycuick 2008). In addition, as temperature decreases with altitude, absolute humidity declines, potentially increasing respiratory water loss (Carmi et al. 1992; Klaassen 1996).

Nevertheless, our results show that saving 20% of flight-incurred energy expenditure during autumn migration through favourable wind selectivity alone could have far-reaching effects for migratory performance and survival. When birds require less energy to complete a migratory segment, they can reduce the number and duration of stopovers needed to replenish fuel stores (Alerstam & Lindström 1990). Fewer stopovers may result in lower exposure to predators and allow individuals to reach wintering grounds earlier and possibly in better body condition, improving access to high-quality resources and competitive opportunities (Alerstam & Lindström 1990). Reduced energetic demand also means birds can carry smaller fat loads. This reduces wing loading (Pennycuick 2008) and cost of transport (Hedenström & Alerstam 1997), thereby making climbing less costly, improving manoeuvrability, and reducing vulnerability to aerial predators (Alerstam & Lindström 1990). Lower energy expenditure further decreases the risk of starvation (Alerstam & Lindström 1990) and reduces the need to use suboptimal stopover sites (Schmaljohann et al. 2022). Collectively, these effects can increase survival during migration. Moreover, arriving at wintering or breeding grounds in better body condition can enhance subsequent reproductive success and overall fitness (Harrison et al. 2011; Nolet et al. 2020; Smith & Moore 2003).

Our findings on wind selectivity and its energetic benefits in short-distance migratory Starlings are particularly intriguing given that flight-related energy expenditure in long-distance migrants can be considerably greater. For long-distance migrants that travel several thousands of kilometres each year and need to traverse ecological barriers along the way, choosing optimal wind conditions may be of even higher importance. For example, in black tailed godwits *Limosa limosa* flying in unfavourable wind conditions during the spring trans-Sahara flights led to increased in-flight mortality rates compared to individuals flying in better wind conditions (Loonstra et al. 2019). Moreover, the reasons behind the recently documented high-altitude flights and frequent altitude changes in multiple long-distance migrants (e.g., Brodersen et al. 2026; Dufour et al. 2025; Sjöberg et al. 2021, 2023) remain poorly understood. The extent to which wind selectivity contributes to such altitude changes and their energetic consequences remains an open question.

While our study allowed analysis of flight behaviour under natural conditions for 113 migratory flights of min 6-hour duration from 30 individuals, several limitations remain. First, stopover locations derived from *GeoPressureR* carry inherent spatial uncertainty, although mostly <100 km (Nussbaumer et al. 2023a), only slightly exceeding ERA5 resolution. Second, we assume linear displacement (“as the crow flies”) between consecutive stopovers since pressure-derived geolocation does not allow precise in-flight location estimation. Although migrants often aim to reduce time aloft (time-minimization theory; Alerstam and Lindström 1990), actual flight paths may still deviate from linear displacement, for example, when following a coastline, avoiding barriers, or navigating along familiar corridors, exposing birds to different wind conditions than estimated. However, large deviations from wind drift are unlikely to go undetected: birds blown off course typically stop to re-orient (Liechti 2006; Richardson 1990b), which would generally be visible in our high-temporal-resolution data, even if very brief stopovers cannot be ruled out entirely. Nevertheless, our study advances over earlier studies that often assumed fixed flight altitudes or relied solely on ground-level or fixed pressure-level wind measurements (Bruderer et al. 1995; Erni et al. 2005; Richardson 1979). Additionally, flight altitudes in our study were obtained from pressure sensor data corrected with ERA5 surface parameters rather than GPS, resulting in low overall uncertainty (mean 10–30 m, maximum 70 m; Nussbaumer & Nussbaumer 2024).

Our results show that in-flight altitude adjustments are closely linked to vertical wind heterogeneity, and high wind selectivity can yield substantial energy savings under headwinds or strong variation among layers. Migrants thus actively navigate the vertical structure of the atmosphere to reduce metabolic costs of flight – a behavioural mechanism with far-reaching consequences for migratory performance, timing, and survival. Our study provides a workable framework to test how widespread in-flight wind selectivity is across species and migratory strategies, enabling quantification of its contribution to annual energy budgets.

## Acknowledgements

We thank Andris Avotiņš for his advice on the analysis of diel-cycle altitude patterns across migratory seasons. We are grateful to D. Spalis, Ā. Ločmele, J. Ločmelis and A. Rimša for their assistance during field work.

## Author contributions

A.P. and M.B. conceived the idea and designed the study. A.P., V.V., O.K., I.D., M.J., V.J., E.M.Z. and M.B. collected the data. A.P., V.V and M.B. analysed the data. R.N. developed *GeoPressureR* and specific functions for wind analysis. A.P. lead the writing of the original draft with substantial contributions from M.B. All authors contributed substantially to revisions.

## Data accessibility statement

The raw and analysed tracking data, including flight altitude profiles, along with wind data used to generate these results https://doi.org/10.5281/zenodo.18467921, as well as the code used for the analyses https://doi.org/10.5281/zenodo.24538322, are deposited on Zenodo Open Repository and will be made open access upon publication.

## References

Able, K.P. (1970). A radar study of the altitude of nocturnal passerine migration. Bird-Band., 41, 282– 290.

Åkesson, S. & Hedenström, A. (2000). Wind selectivity of migratory flight departures in birds. Behav. Ecol. Sociobiol., 47, 140–144.

Alerstam, T. & Lindström, Å. (1990). Optimal bird migration: the relative importance of time, energy, and safety. In: Bird migration: physiology and ecophysiology (ed. Gwinner, E.). Springer, Berlin, pp. 331–351.

Barron, D.G., Brawn, J.D. & Weatherhead, P.J. (2010). Meta-analysis of transmitter effects on avian behaviour and ecology. Methods Ecol. Evol., 1, 180–187.

Battley, P. (1997). The Northward Migration of Arctic Waders in New Zealand: Departure Behaviour, Timing and Possible Migration Routes of Red Knots and Bar-tailed Godwits from Farewell Spit, North-West Nelson. Emu - Austral Ornithol., 97, 108–120.

Berthold, P. (2001). Bird migration: a general survey. 2nd edn. Oxford University Press, Oxford.

Bowlin, M.S., Enstrom, D.A., Murphy, B.J., Plaza, E., Jurich, P. & Cochran, J. (2015). Unexplained altitude changes in a migrating thrush: long-flight altitude data from radio-telemetry. Auk Ornithol. Adv., 132, 808–816.

Brlík, V., Koleček, J., Burgess, M., Hahn, S., Humple, D., Krist, M., et al. (2020). Weak effects of geolocators on small birds: A meta-analysis controlled for phylogeny and publication bias. J. Anim. Ecol., 89, 207–220.

Brodersen, J.E., Lindström, Å., Alerstam, T., Andersson, A., Bäckman, J., Hedenström, A., et al. (2026). Low cost of shifting flight altitude in migrating great snipes. J. Exp. Biol., 229, jeb252022.

Bruderer, B. & Boldt, A. (2001). Flight characteristics of birds: I. Radar measurements of speeds. Ibis, 143, 178–204.

Bruderer, B., Peter, D., Boldt, A. & Liechti, F. (2010). Wing-beat characteristics of birds recorded with tracking radar and cine camera. Ibis, 152, 272–291.

Bruderer, B., Underhill, L. & Liechti, F. (1995). Altitude choice by night migrants in a desert area predicted by meteorological factors. Ibis, 137, 44–55.

Carey, C. (Ed.). (1996). Avian Energetics and Nutritional Ecology. Springer US, Boston, MA.

Carmi, N., Pinshow, B., Porter, W.P. & Jaeger, J. (1992). Water and energy limitations on flight duration in small migrating birds. The Auk, 109, 268–276.

Dokter, A.M., Shamoun-Baranes, J., Kemp, M.U., Tijm, S. & Holleman, I. (2013). High Altitude Bird Migration at Temperate Latitudes: A Synoptic Perspective on Wind Assistance. PLoS ONE, 8, e52300.

Dufour, P., Nussbaumer, R., Briedis, M., Bocher, P., Conway, G., Coulomb, Y., et al. (2025). Ecological barrier crossing strategies in small migratory birds depend on wing morphology and plumage color. iScience, 29, 114466.

Erni, B., Liechti, F. & Bruderer, B. (2005). The role of wind in passerine autumn migration between Europe and Africa. Behav. Ecol., 16, 732–740.

Gauthreaux, S.A.J. (1991). The flight behavior of migrating birds in changing wind fields: radar and visual analyses. Am. Zool., 31, 187–204.

Gill, R.E., Jr, Tibbitts, T.L., Douglas, D.C., Handel, C.M., Mulcahy, D.M., Gottschalck, J.C., et al. (2009). Extreme endurance flights by landbirds crossing the Pacific Ocean: ecological corridor rather than barrier? Proc. R. Soc. B Biol. Sci., 276, 447–457.

Guglielmo, C.G. (2018). Obese super athletes: fat-fueled migration in birds and bats. J. Exp. Biol., 221, jeb165753.

Harris, S.M., Bishop, C.M., Bond, S., Fernandes, P.G., Guilford, T., Lewin, P.J., et al. (2025). Adjustable wind selectivity in shearwaters implies knowledge of the foraging landscape. Curr. Biol., 35, 889–897.

Harrison, X.A., Blount, J.D., Inger, R., Norris, D.R. & Bearhop, S. (2011). Carry-over effects as drivers of fitness differences in animals. J. Anim. Ecol., 80, 4–18.

Hedenström, A. & Alerstam, T. (1997). Optimum Fuel Loads in Migratory Birds: Distinguishing Between Time and Energy Minimization. J. Theor. Biol., 189, 227–234.

Hersbach, H., Bell, B., Berrisford, P., Biavati, G., Horányi, A., Muñoz Sabater, J., et al. (2023). ERA5 hourly data on pressure levels from 1940 to present. Available at: 10.24381/cds.bd0915c6. Last accessed 14 October 2026.

Horton, K.G., Van Doren, B.M., Stepanian, P.M., Farnsworth, A. & Kelly, J.F. (2016). Where in the air? Aerial habitat use of nocturnally migrating birds. Biol. Lett., 12, 20160591.

Hurrell, J.W. & Deser, C. (2010). North Atlantic climate variability: The role of the North Atlantic Oscillation. Impact Clim. Var. Mar. Ecosyst. Comp. Approach, 79, 231–244.

Kemp, M.U., Shamoun-Baranes, J., Dokter, A.M., van Loon, E. & Bouten, W. (2013). The influence of weather on the flight altitude of nocturnal migrants in mid-latitudes. Ibis, 155, 734–749.

Klaassen, M. (1996). Metabolic Constraints on Long-Distance Migration in Birds. J. Exp. Biol., 199, 57– 64.

Klein Heerenbrink, M. (2023). afpt: Tools for Modelling of Animal Flight Performance. version 1.1.0.4. Available at: https://github.com/marcoklh/afpt-r.

Liechti, F. (2006). Birds: blowin’ by the wind? J. Ornithol., 147, 202–211.

Liechti, F., Bauer, S., Dhanjal-Adams, K.L., Emmenegger, T., Zehtindjiev, P. & Hahn, S. (2018). Miniaturized multi-sensor loggers provide new insight into year-round flight behaviour of small trans-Sahara avian migrants. Mov. Ecol., 6, 19.

Liechti, F. & Schmaljohann, H. (2007). Wind-governed flight altitudes of nocturnal spring migrants over the Sahara. Ostrich-J. Afr. Ornithol., 78, 337–341.

Lindström, Å., Alerstam, T., Andersson, A., Bäckman, J., Bahlenberg, P., Bom, R., et al. (2021). Extreme altitude changes between night and day during marathon flights of great snipes. Curr. Biol., 31, 3433–3439.

Mateos, M. & Liechti, F. (2012). How do diurnal long-distance migrants select flight altitude in relation to wind? Behav. Ecol., 23, 403–409.

McWilliams, S.R., Guglielmo, C., Pierce, B. & Klaassen, M. (2004). Flying, fasting, and feeding in birds during migration: a nutritional and physiological ecology perspective. J. Avian Biol., 35, 377– 393.

Muñoz Sabater, J. (2019). ERA5-Land hourly data from 1950 to present. Available at: 10.24381/cds.e2161bac. Last accessed 14 October 2026.

Newton, I. (2023). The migration ecology of birds. Academic Press, London.

Nolet, B.A., Schreven, K.H.T., Boom, M.P. & Lameris, T.K. (2020). Contrasting effects of the onset of spring on reproductive success of Arctic-nesting geese. The Auk, 137, 1–9.

Norevik, G., Åkesson, S., Andersson, A., Bäckman, J. & Hedenström, A. (2021). Flight altitude dynamics of migrating European nightjars across regions and seasons. J. Exp. Biol., 224, jeb242836.

Nudds, R.L. & Bryant, D.M. (2000). The energetic cost of short flights in birds. J. Exp. Biol., 203, 1561– 1572.

Nussbaumer, R. & Gravey, M. (2024). GeoPressureR: Global Positioning by Atmospheric Pressure. version v3.2.0. Available at: 10.5281/zenodo.10642935.

Nussbaumer, R., Gravey, M., Briedis, M. & Liechti, F. (2023a). Global positioning with animal-borne pressure sensors. Methods Ecol. Evol., 14, 1104–1117.

Nussbaumer, R., Gravey, M., Briedis, M. & Liechti, F. (2023b). Reconstructing bird trajectories from pressure and wind data using a highly optimized hidden Markov model. Methods Ecol. Evol., 14, 1118–1129.

Nussbaumer, R. & Nussbaumer, A. (2024). GeoPressureManual: User Manual for GeoPressureR. Available at: 10.5281/zenodo.10799355.

Pennycuick, C.J. (1969). The mechanics of bird migration. Ibis, 111, 525–556. Pennycuick, C.J. (2008). Modelling the flying bird. Elsevier, London.

Pyle, P., Nur, N., Henderson, R. & Desante, D. (1993). The Effects of Weather and Lunar Cycle on Nocturnal Migration of Landbirds at Southeast Farallon Island, California. The Condor, 95, 343–361.

R Core Team. (2024). R: A Language and Environment for Statistical Computing. R Foundation for Statistical Computing, Vienna, Austria.

Richardson, W.J. (1979). Southeastward shorebird migration over Nova Scotia and New Brunswick in autumn: a radar study. Can. J. Zool., 57, 107–124.

Richardson, W.J. (1990a). Timing of Bird Migration in Relation to Weather: Updated Review. In: Bird Migration (ed. Gwinner, E.). Springer Berlin Heidelberg, Berlin, Heidelberg, pp. 78–101.

Richardson, W.J. (1990b). Wind and orientation of migrating birds: A review. Experientia, 46, 416– 425.

Scacco, M., Smaers, J.B., Werber, Y., Aghababyan, K., Arrondo, E., Aschenborn, O., et al. (2026). Airflows redefine the cost of aerial transport in the wild. Science, 394, 347–352.

Schaub, M., Liechti, F. & Jenni, L. (2004). Departure of migrating European robins, Erithacus rubecula, from a stopover site in relation to wind and rain. Anim. Behav., 67, 229–237.

Schmaljohann, H., Eikenaar, C. & Sapir, N. (2022). Understanding the ecological and evolutionary function of stopover in migrating birds. Biol. Rev., 97, 1231–1252.

Senner, N.R., Stager, M., Verhoeven, M.A., Cheviron, Z.A., Piersma, T. & Bouten, W. (2018). High-altitude shorebird migration in the absence of topographical barriers: avoiding high air temperatures and searching for profitable winds. Proc. R. Soc. B Biol. Sci., 285, 20180569.

Sjöberg, S., Andersson, A., Bäckman, J., Hansson, B., Malmiga, G., Tarka, M., et al. (2023). Solar heating may explain extreme diel flight altitude changes in migrating birds. Curr. Biol., 33, 4232–4237.e2.

Sjöberg, S., Malmiga, G., Nord, A., Andersson, A., Bäckman, J., Tarka, M., et al. (2021). Extreme altitudes during diurnal flights in a nocturnal songbird migrant. Science, 372, 646–648.

Smith, R. & Moore, F. (2003). Arrival fat and reproductive performance in a long-distance passerine migrant. Oecologia, 134, 325–331.

Stull, R.B. (2015). Practical meteorology: an algebra-based survey of atmospheric science. University of British Columbia, Department of Earth, Ocean and Atmospheric Sciences, Vancouver.

Tobalske, B.W. (1995). Neuromuscular control and kinematics of intermittent flight in the European starling (Sturnus vulgaris). J. Exp. Biol., 198 Pt 6, 1259–1273.

Vīgants, V., Keišs, O., Dinsbergs, I., Jaunzemis, V., Zacmane, E.M., Priedniece, A., et al. (2023). Migration strategies, performance and annual activity budget in a short-distance migrant, the common starling Sturnus vulgaris. J. Avian Biol., 2023, e03080.

Winkler, D.W., Shamoun-Baranes, J. & Piersma, T. (2016). Avian migration and dispersal. In: Handbook of Bird Biology (eds. Lovette, I.J. & Fitzpatrick, J.W.). John Wiley & Sons, Chichester, pp. 453– 492.

Wood, S., Scheipl, F. & Schneuwly, A. (2026). gamm4: Generalized Additive Mixed Models using “mgcv” and “lme4.” version 0.3–0. Available at: https://CRAN.R-project.org/package=gamm4.

Wood, S.N. (2017). Generalized additive models: an introduction with R. 2nd edn. Chapman and Hall/CRC, Boca Raton.

