## Supporting information for "Vertical in-flight wind selectivity enables substantial energy savings in a migratory bird"

**Figure S1.** Variation in flight altitudes and their corresponding wind conditions recorded for individual Starling during flights of minimum 6 hours. On each panel, background colours indicate the strength of wind support at different altitudes during the flight, where negative values correspond to headwinds and positive to tailwinds. The black solid line depicts the recorded flight path, and the doted light indicates ideal altitude path to maximize windsupport. The dark polygon on the bottom of the panels denotes the average ground elevation.

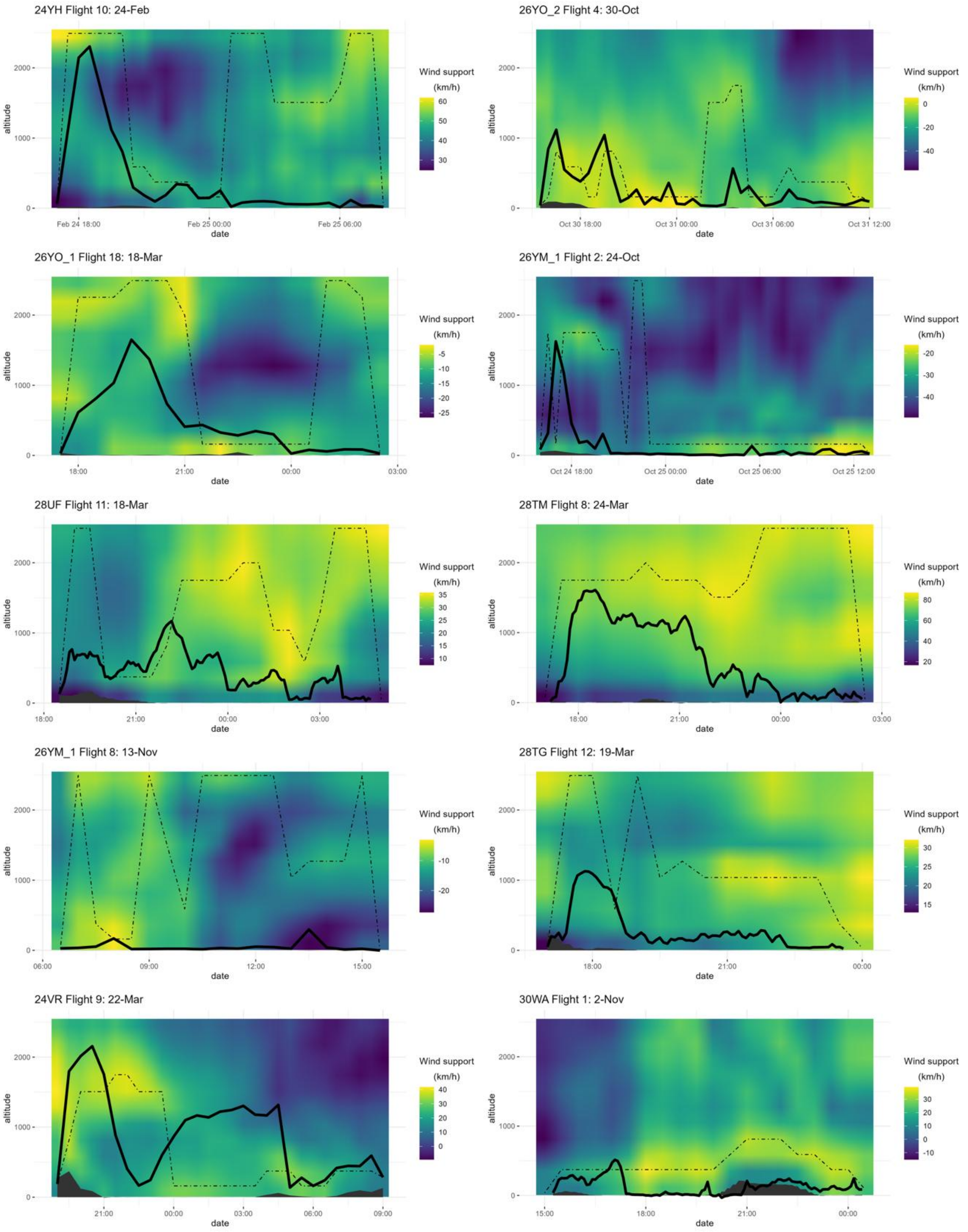

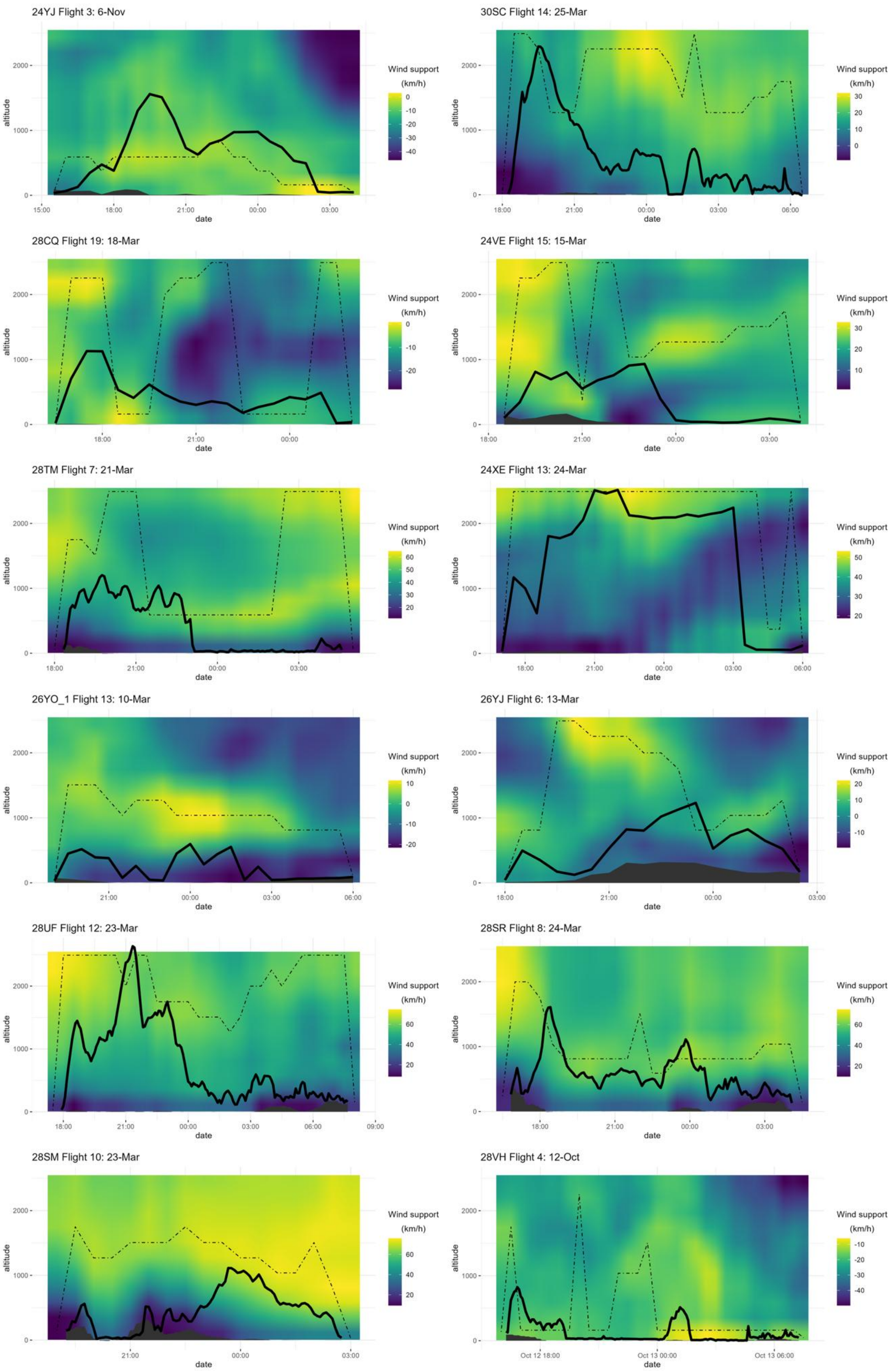

24YJ Flight 3: 6-Nov
altitude
date
Wind support (km/h)
0
-10
-20
-30
-40
15:00
18:00
21:00
00:00
03:00
30SC Flight 14: 25-Mar
Wind support (km/h)
30
20
10
0
18:00
21:00
00:00
03:00
06:00
28CQ Flight 19: 18-Mar
Wind support (km/h)
0
-10
-20
24VE Flight 15: 15-Mar
Wind support (km/h)
30
20
10
28TM Flight 7: 21-Mar
Wind support (km/h)
60
50
40
30
20
24XE Flight 13: 24-Mar
Wind support (km/h)
50
40
30
20
26YO_1 Flight 13: 10-Mar
Wind support (km/h)
10
0
-10
-20
26YJ Flight 6: 13-Mar
Wind support (km/h)
20
10
0
-10
28UF Flight 12: 23-Mar
Wind support (km/h)
60
40
20
09:00
28SR Flight 8: 24-Mar
Wind support (km/h)
60
40
20
28SM Flight 10: 23-Mar
Wind support (km/h)
60
40
20
28VH Flight 4: 12-Oct
Wind support (km/h)
-10
-20
-30
-40
Oct 12 18:00
Oct 13 00:00
Oct 13 06:00
2000
1000
0

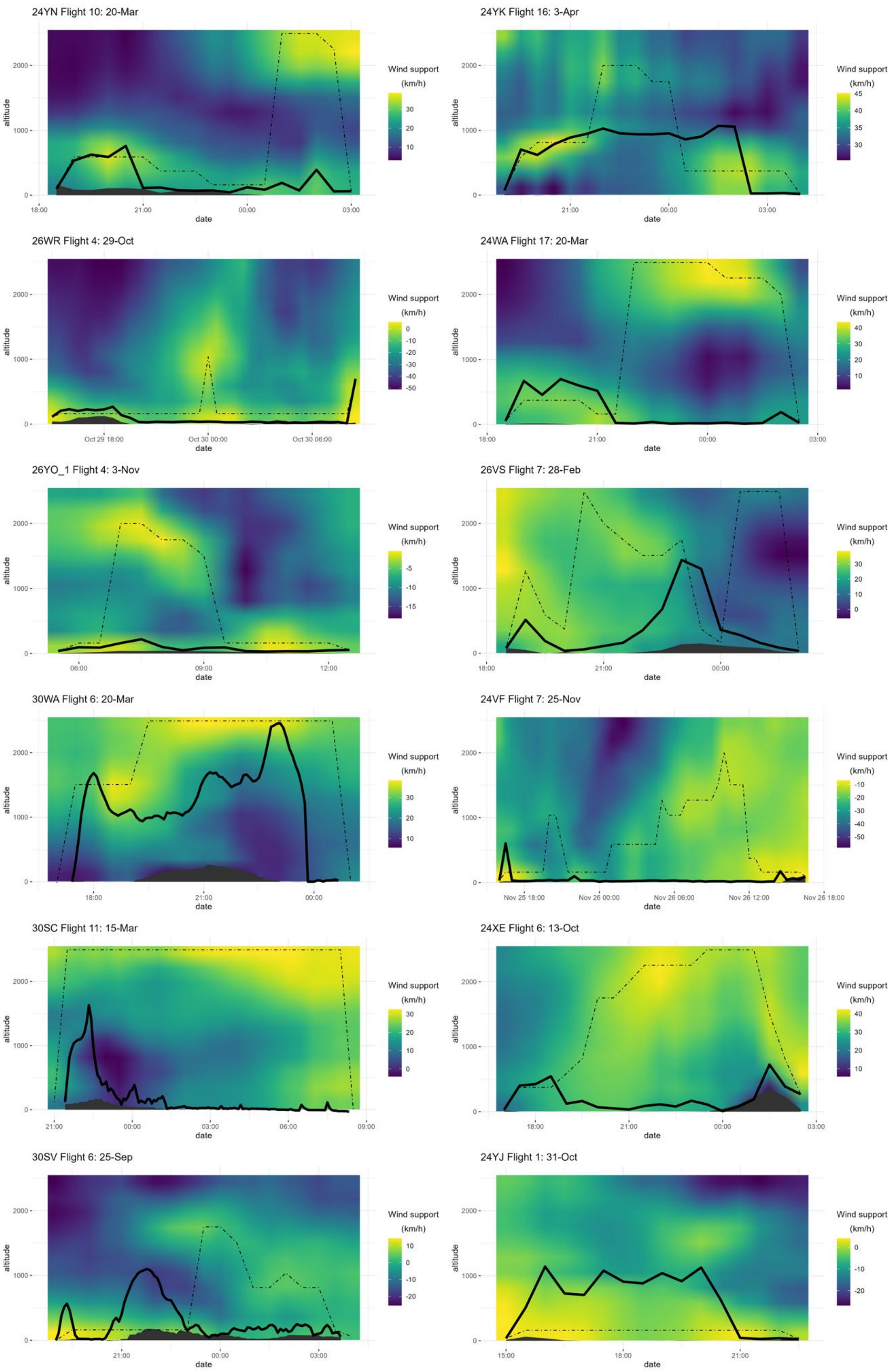
24YN Flight 10: 20-Mar
24YK Flight 16: 3-Apr
26WR Flight 4: 29-Oct
24WA Flight 17: 20-Mar
26YO_1 Flight 4: 3-Nov
26VS Flight 7: 28-Feb
30WA Flight 6: 20-Mar
24VF Flight 7: 25-Nov
30SC Flight 11: 15-Mar
24XE Flight 6: 13-Oct
30SV Flight 6: 25-Sep
24YJ Flight 1: 31-Oct
altitude
date
Wind support (km/h)

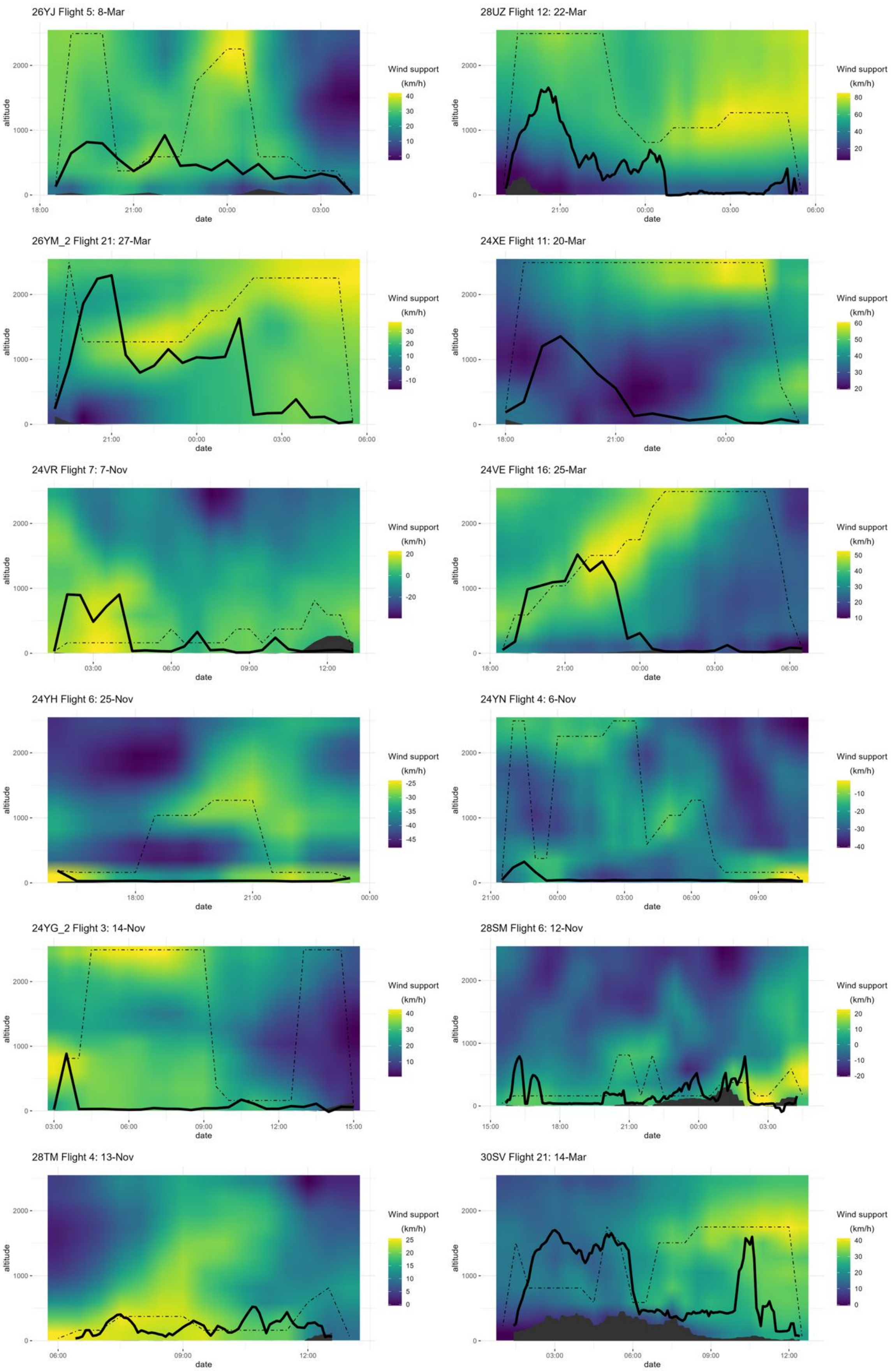
26YJ Flight 5: 8-Mar
28UZ Flight 12: 22-Mar
26YM_2 Flight 21: 27-Mar
24XE Flight 11: 20-Mar
24VR Flight 7: 7-Nov
24VE Flight 16: 25-Mar
24YH Flight 6: 25-Nov
24YN Flight 4: 6-Nov
24YG_2 Flight 3: 14-Nov
28SM Flight 6: 12-Nov
28TM Flight 4: 13-Nov
30SV Flight 21: 14-Mar
altitude
date
Wind support
(km/h)

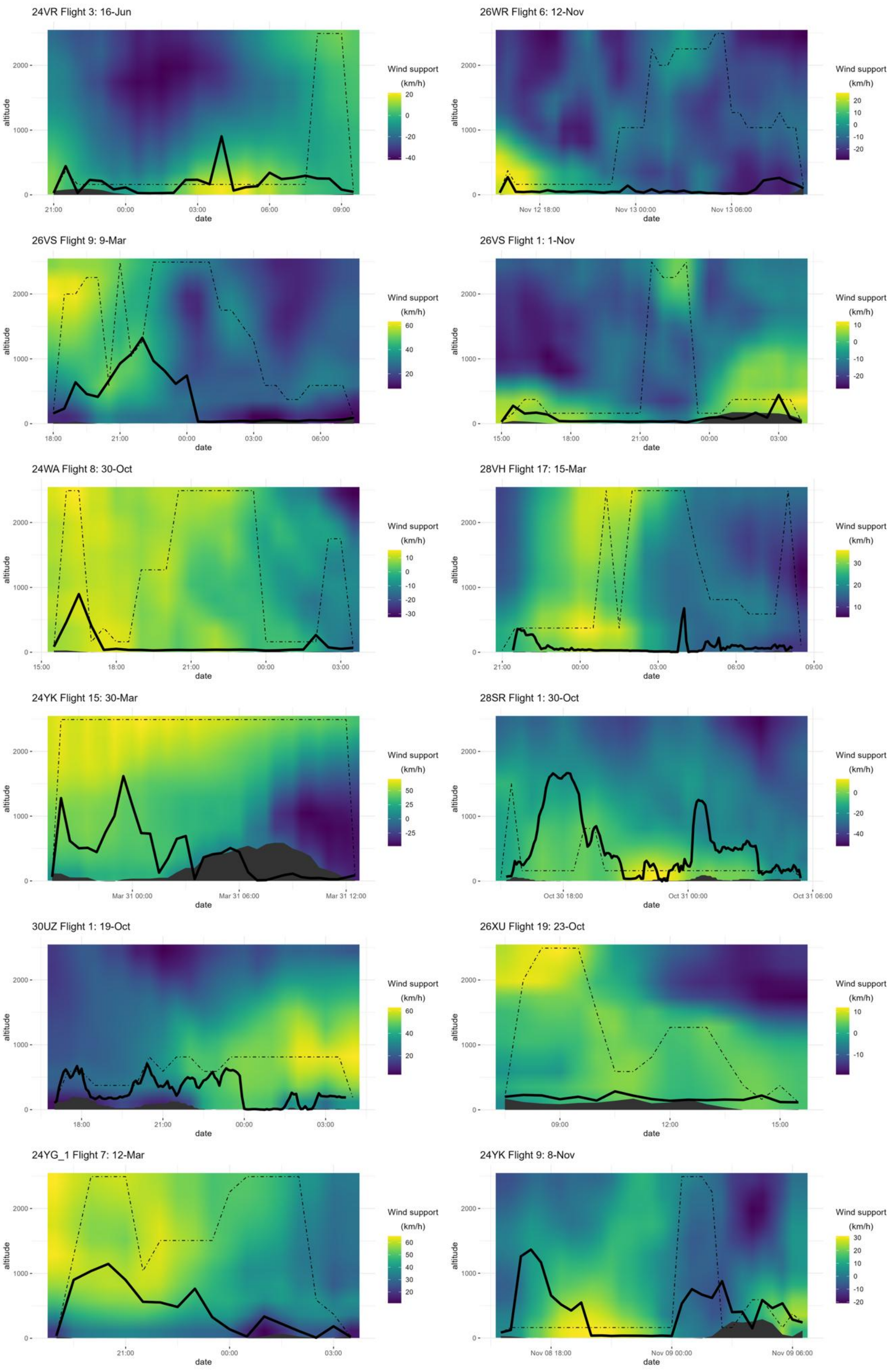
24VR Flight 3: 16-Jun
26WR Flight 6: 12-Nov
26VS Flight 9: 9-Mar
26VS Flight 1: 1-Nov
24WA Flight 8: 30-Oct
28VH Flight 17: 15-Mar
24YK Flight 15: 30-Mar
28SR Flight 1: 30-Oct
30UZ Flight 1: 19-Oct
26XU Flight 19: 23-Oct
24YG_1 Flight 7: 12-Mar
24YK Flight 9: 8-Nov
altitude
date
Wind support
(km/h)

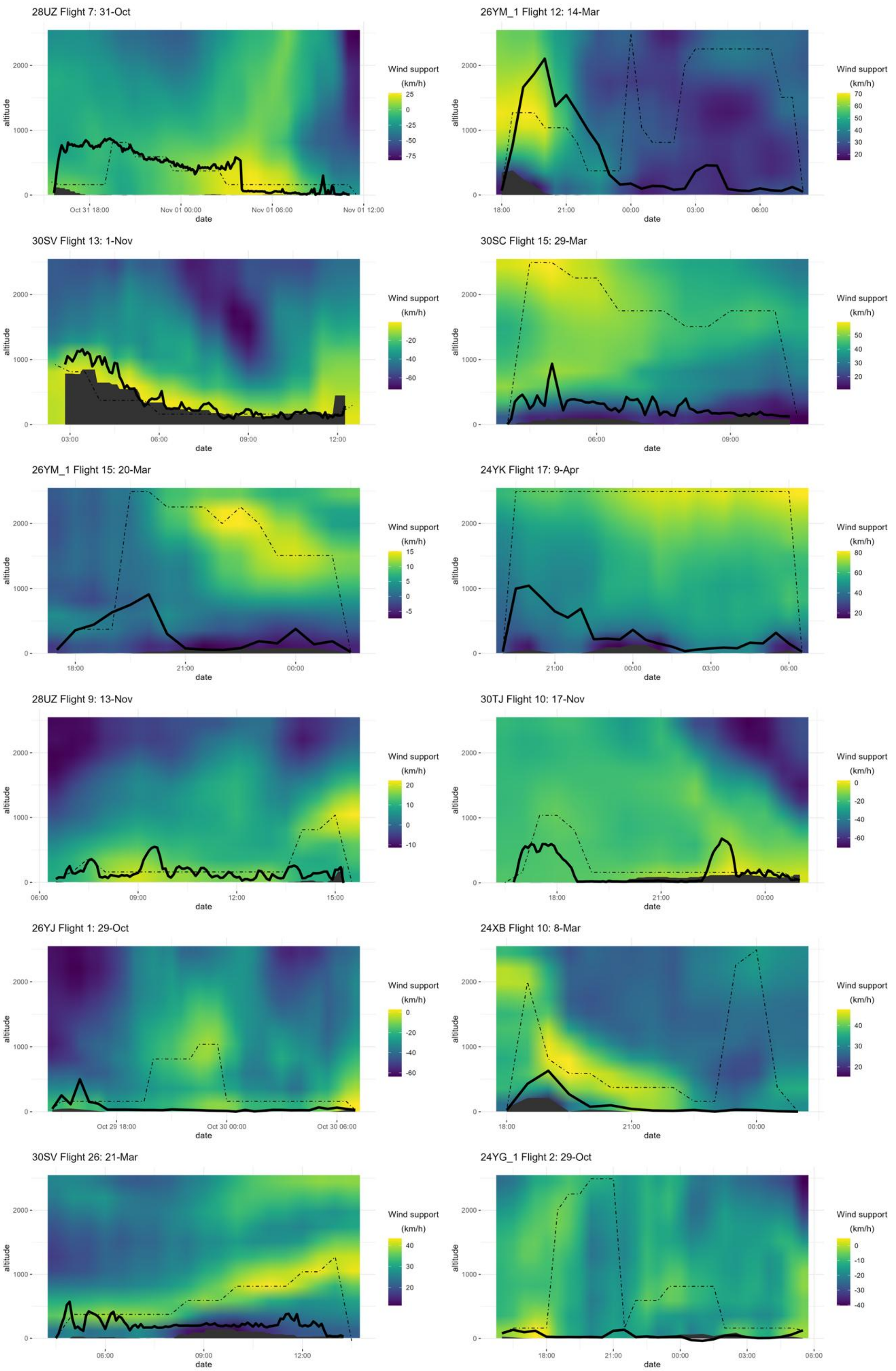
28UZ Flight 7: 31-Oct
26YM_1 Flight 12: 14-Mar
30SV Flight 13: 1-Nov
30SC Flight 15: 29-Mar
26YM_1 Flight 15: 20-Mar
24YK Flight 17: 9-Apr
28UZ Flight 9: 13-Nov
30TJ Flight 10: 17-Nov
26YJ Flight 1: 29-Oct
24XB Flight 10: 8-Mar
30SV Flight 26: 21-Mar
24YG_1 Flight 2: 29-Oct
Wind support (km/h)
altitude
date

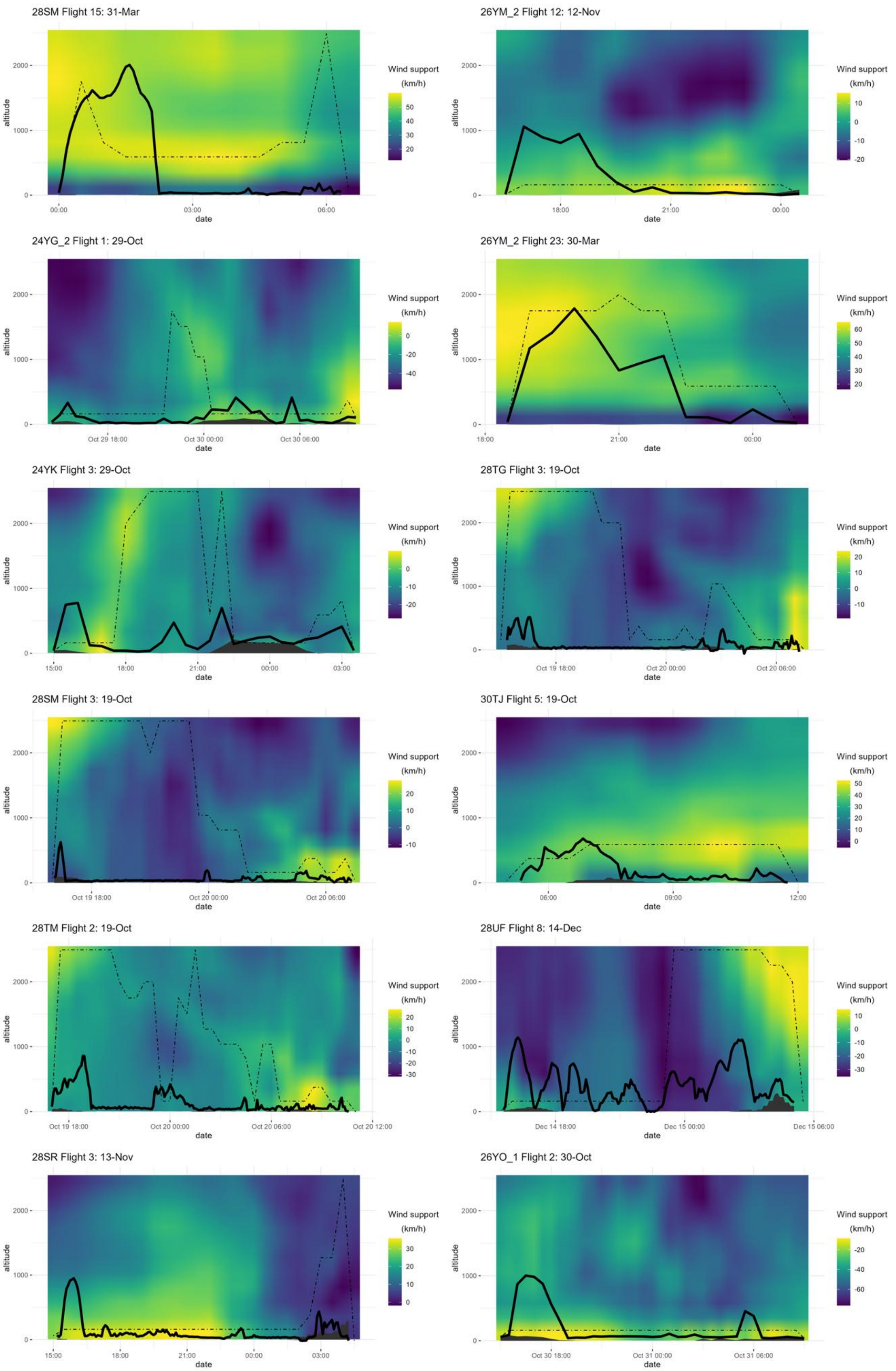
28SM Flight 15: 31-Mar
26YM_2 Flight 12: 12-Nov
24YG_2 Flight 1: 29-Oct
26YM_2 Flight 23: 30-Mar
24YK Flight 3: 29-Oct
28TG Flight 3: 19-Oct
28SM Flight 3: 19-Oct
30TJ Flight 5: 19-Oct
28TM Flight 2: 19-Oct
28UF Flight 8: 14-Dec
28SR Flight 3: 13-Nov
26YO_1 Flight 2: 30-Oct
altitude
date
Wind support
(km/h)

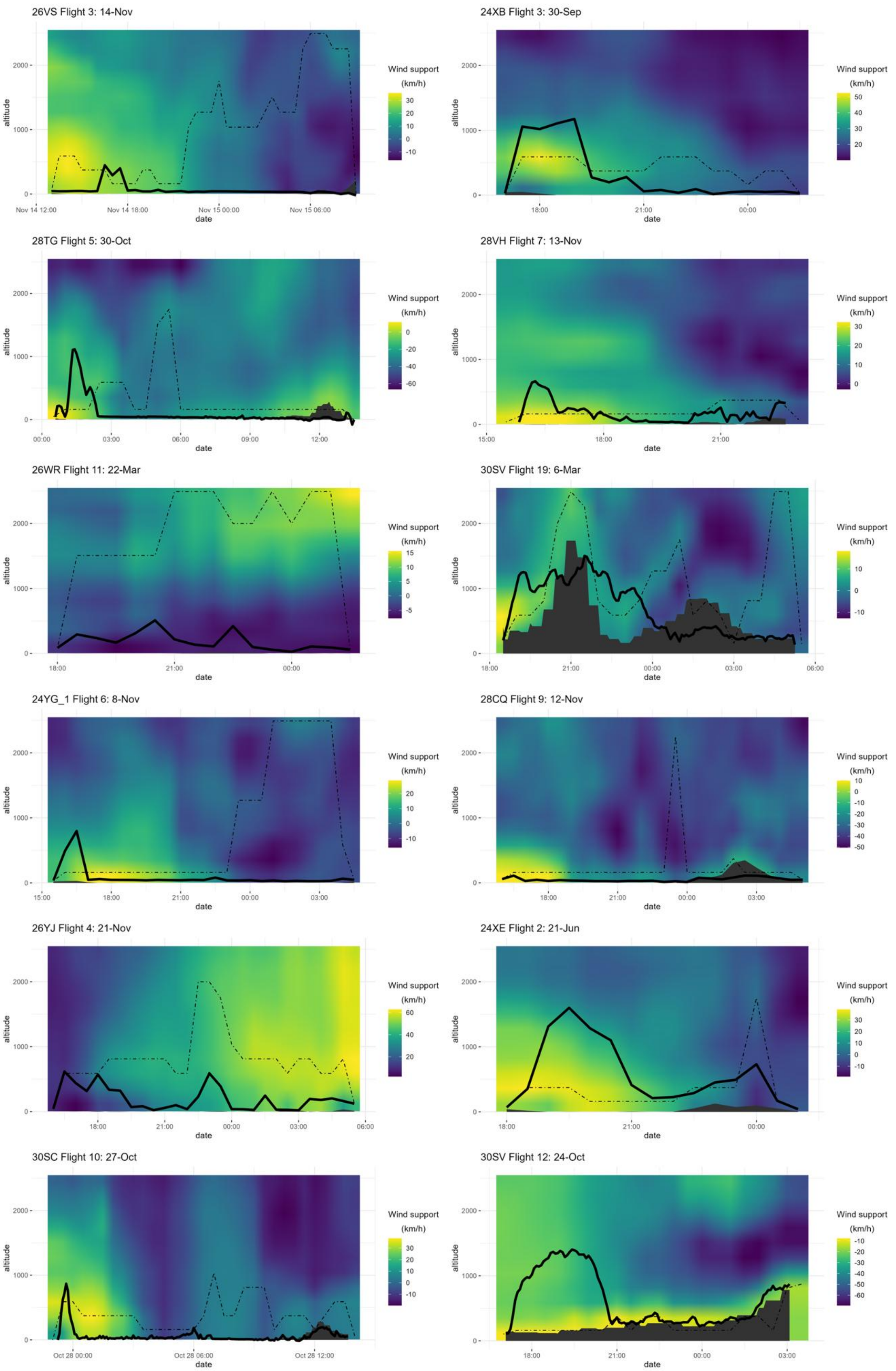
26VS Flight 3: 14-Nov
24XB Flight 3: 30-Sep
28TG Flight 5: 30-Oct
28VH Flight 7: 13-Nov
26WR Flight 11: 22-Mar
30SV Flight 19: 6-Mar
24YG_1 Flight 6: 8-Nov
28CQ Flight 9: 12-Nov
26YJ Flight 4: 21-Nov
24XE Flight 2: 21-Jun
30SC Flight 10: 27-Oct
30SV Flight 12: 24-Oct
Wind support (km/h)
altitude
date

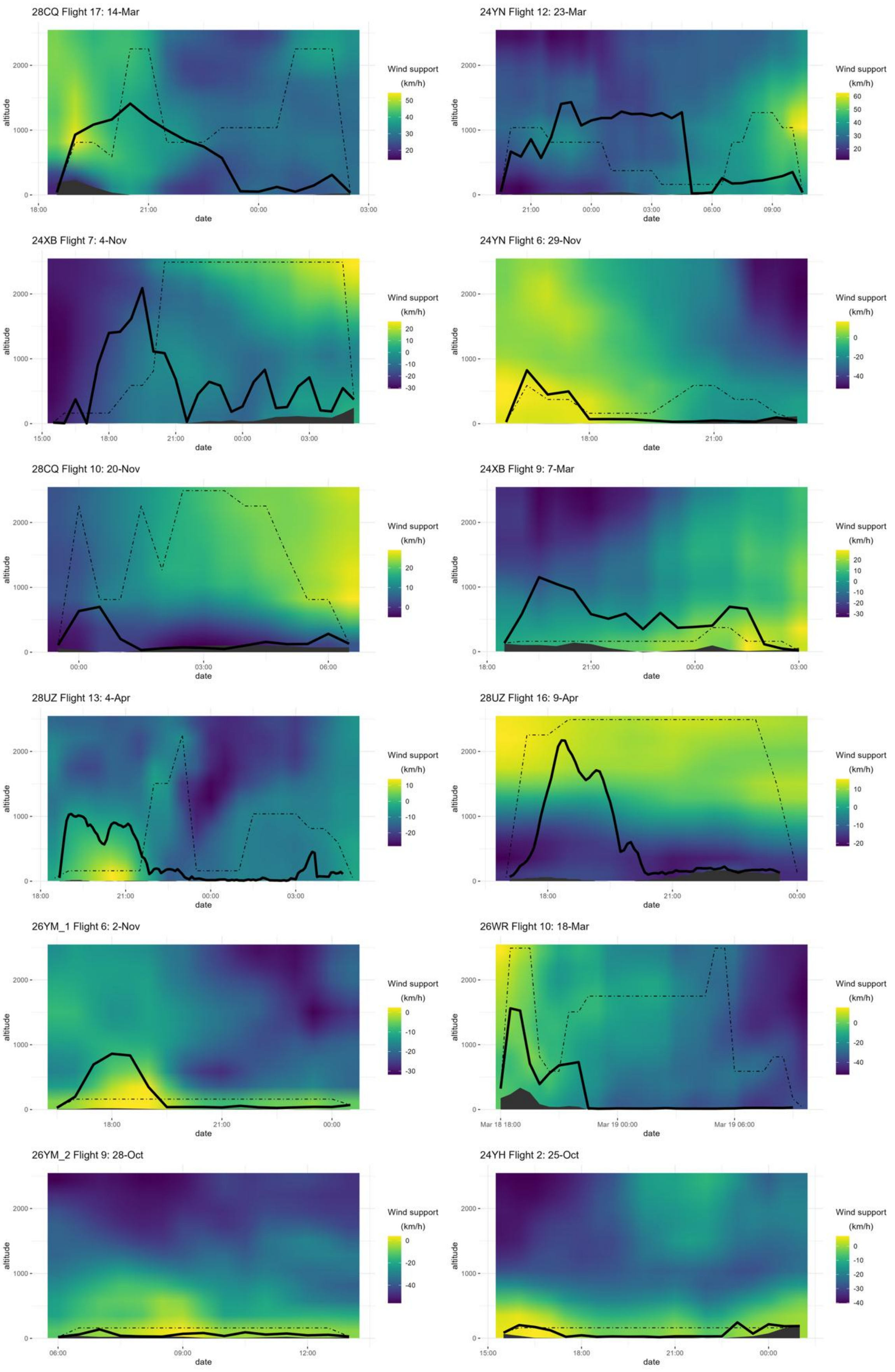

28CQ Flight 17: 14-Mar
24YN Flight 12: 23-Mar
24XB Flight 7: 4-Nov
24YN Flight 6: 29-Nov
28CQ Flight 10: 20-Nov
24XB Flight 9: 7-Mar
28UZ Flight 13: 4-Apr
28UZ Flight 16: 9-Apr
26YM_1 Flight 6: 2-Nov
26WR Flight 10: 18-Mar
26YM_2 Flight 9: 28-Oct
24YH Flight 2: 25-Oct
Wind support (km/h)
altitude
date
Mar 18 18:00
Mar 19 00:00
Mar 19 06:00

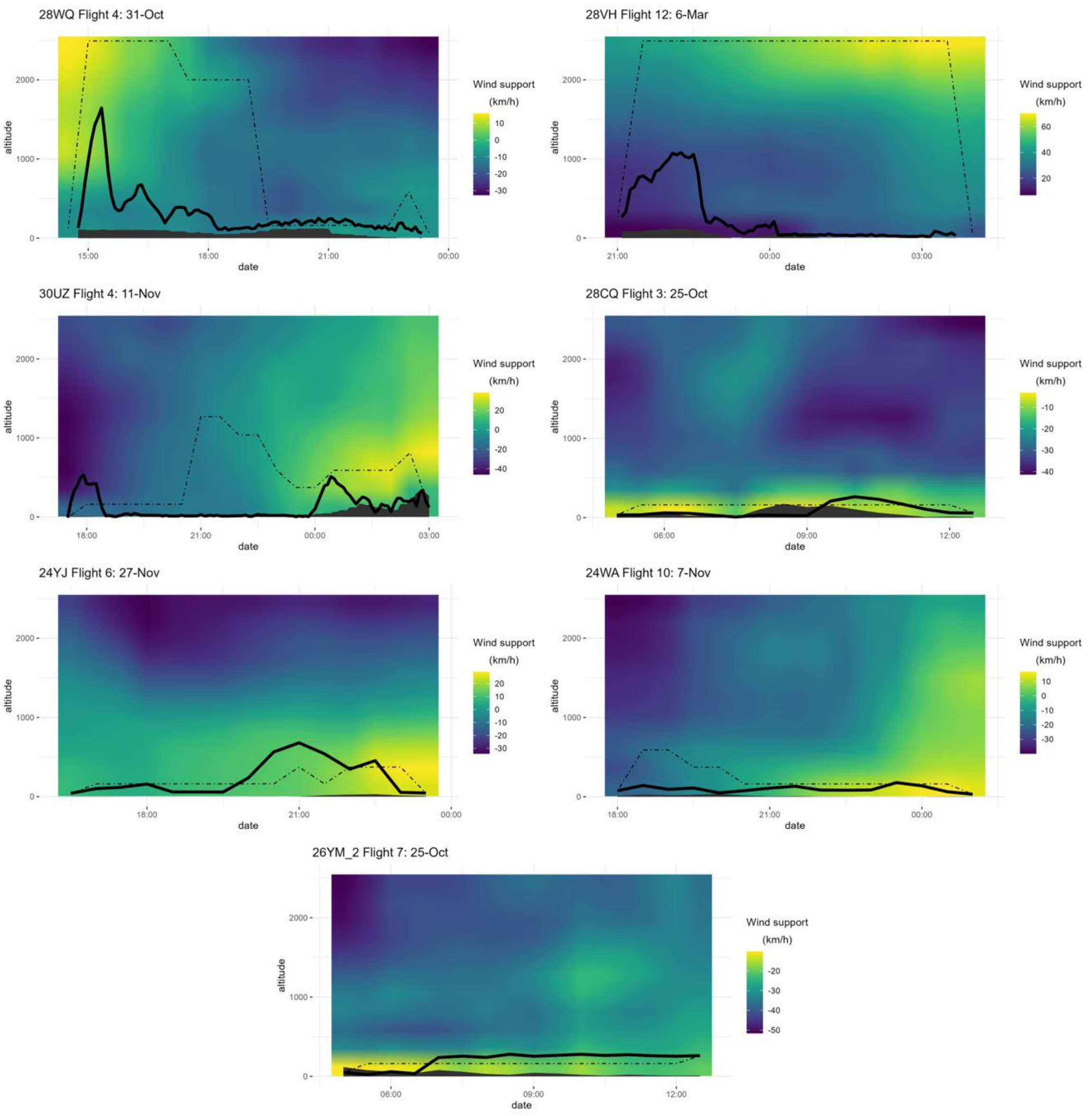
28WQ Flight 4: 31-Oct
altitude
0
1000
2000
date
15:00
18:00
21:00
00:00
Wind support
(km/h)
10
0
-10
-20
-30
28VH Flight 12: 6-Mar
altitude
0
1000
2000
date
21:00
00:00
03:00
Wind support
(km/h)
60
40
20
30UZ Flight 4: 11-Nov
altitude
0
1000
2000
date
18:00
21:00
00:00
03:00
Wind support
(km/h)
20
0
-20
-40
28CQ Flight 3: 25-Oct
altitude
0
1000
2000
date
06:00
09:00
12:00
Wind support
(km/h)
-10
-20
-30
-40
24YJ Flight 6: 27-Nov
altitude
0
1000
2000
date
18:00
21:00
00:00
Wind support
(km/h)
20
10
0
-10
-20
-30
24WA Flight 10: 7-Nov
altitude
0
1000
2000
date
18:00
21:00
00:00
Wind support
(km/h)
10
0
-10
-20
-30
26YM_2 Flight 7: 25-Oct
altitude
0
1000
2000
date
06:00
09:00
12:00
Wind support
(km/h)
-20
-30
-40
-50
